# Recovery of silver fir (*Abies alba* Mill.) seedlings from ungulate browsing mirrors soil nitrogen availability

**DOI:** 10.1101/2021.04.12.439531

**Authors:** Katalin Csilléry, Nina Buchmann, Oliver Brendel, Arthur Gessler, Alexandra Glauser, Andrea Doris Kupferschmid

## Abstract

*Abies alba* has a high potential for mitigating climate change in European mountain forests, yet, its natural regeneration is severely limited by ungulate browsing. Here, we simulated browsing in a common garden experiment to study growth and physiological traits, measured from bulk needles, using a randomized block design with two levels of browsing severity and seedlings originating from 19 populations across Switzerland. Genetic factors explained most variation in growth (on average, 51.5%) and physiological traits (10.2%) under control conditions, while heavy browsing considerably reduced the genetic effects on growth (to 30%), but doubled those on physiological traits related to C storage. While browsing reduced seedling height, it also lowered seedlings’ water use efficiency (decreased *δ*^13^C) and N supply by mycorrhizal fungi as indicated by an increase in *δ*^15^N. Different populations reacted differently to browsing stress, and for Height, Starch and *δ*^15^N, population differences appeared to be the result of natural selection. We found that the fastest growing populations, originating from the warmest regions, decreased their needle starch level the most as a reaction to heavy browsing, suggesting a potential genetic underpinning for a growth-storage trade-off. Further, we found that seedlings originating from mountain populations growing on steep slopes had a significantly lower N discrimination in the common garden than those originating from flat areas, indicating that they have been selected to grow on N poor, potentially drained, soils. This finding was corroborated by the fact that N concentration in adult needles was lower on steep slopes than on flat ground, strongly indicating that steep slopes are the most N poor environments. Seedlings from these poor environments generally had a low growth rate and high storage, thus might be slower to recover from browsing stress than fast growing provenances from the warm environments with developed soils, such as the Swiss plateau.

## Introduction

The forested land area in Europe has increased by 56% over the past 100 years (Fuchs *et al*., 2015). Although climate change is partly responsible for this area gain, reforestation is also associated with management activities, such as the abandonment of agricultural land and afforestation efforts to increase timber volume and economic benefit (Seidl *et al*., 2011). Ungulates spontaneously recolonized these new habitats and were further aided by artificial re-introductions. Further, until recently, due to the lack of natural predators and a decrease in hunting for big games, ungulate numbers kept increasing (Apollonio *et al*., 2010). As a result, ungulate browsing is a major driver of forest succession in the northern hemisphere and challenges the establishment of future tree generations (Tanentzap *et al*., 2009; Apollonio *et al*., 2017). Since ungulates selectively browse certain tree species, such damage can have long lasting impact on the forest species composition (Klopčič *et al*., 2017; Ramirez *et al*., 2019).

The most common effects of ungulate browsing are the removal of buds, thus apical meristem tissue and the removal of shoots, which reduces the photosynthesizing leaf area and changes the root-to-shoot ratio (Hoogesteger & Karlsson, 1992; Rhodes & Clair, 2018). Trees are often found to recover well from light to moderate browsing stress via different compensatory mechanisms at the morphological level such as increased leaf size (Lehtilä *et al*., 2000), overcompensated growth in the leading bud (O’Reilly-Wapstra *et al*., 2014), growing side shoots (Kupferschmid & Heiri, 2019), but also in their reproductive strategy such as increased production of female strobili (Allison, 1990). However, browsing can also severely limit growth and cause seedling mortality (Kupferschmid, 2017; Rhodes & Clair, 2018). Browsing damage and recovery depend on several factors including the intensity and timing of browsing, but also on the stress status and the ontogenetic stage of the tree (Kupferschmid, 2017). Browsing stress generally affects the most severely the early life-stages, which are already the most sensitive to environmental fluctuations and climate change related risks (Talluto *et al*., 2017).

Early studies argued that browsing stress causes carbon (C) limitation in agreement with the C-supply-centered view of tree growth (Ericsson *et al*., 1980; Chapin III *et al*., 1980). However, this view has largely been challenged during the past two decades. The emerging picture is that trees are rarely, if at all, C limited (Körner, 2003; Millard & Grelet, 2010; Sala *et al*., 2012; McDowell *et al*., 2008). For example, trees have been observed to accumulate large amounts of non-structural carbohydrates (NSC) even in the presence of factors that limit their photosynthesis, such as under severe water deficit (Bréda *et al*., 2006), defoliation (Wiley *et al*., 2013), or light limitation (Weber *et al*., 2018). Although seasonal fluctuations in NSC levels have been observed in seedlings, such as during bud burst or recovery from herbivory (Gill, 1992), there was no evidence for C limitation *per se* (Palacio *et al*., 2008).

In contrast to C, nitrogen (N) is stored and seasonally remobilized; for example, N storage pools disappear after bud flushing (Millard & Grelet, 2010). In conifers, N is mainly present in young needles as photosynthetic proteins, as RuBisCo (Millard *et al*., 2001; Camm, 1993) or as amino acids (Schneider *et al*., 1996). Thus, ungulate browsing can drastically reduce N pools, and the recovery from browsing depends on seedlings’ capacity to remobilize N from other tissue and on their N acquisition from the environment. Conifers commonly grow in nutrient-poor sites because they can store N in their foliage, which can be easily mobilized, thus, their reserves are particularly susceptible to loss by herbivory (Millard & Grelet, 2010). Most root N uptake occurs via biotic interactions in the rhizosphere such as mycorrhizal symbiosis, associations with free-living fungi and bacteria, and endophytic bacteria that can increase the efficiency of N acquisition and assimilation (Millard & Grelet, 2010). The stable N isotope composition (*δ*^15^N) of plant tissues is commonly used to identify variations in N supply and uptake. *δ*^15^N is determined by the relative contribution of external N source with different *δ*^15^N (e.g. different soil pools, atmosphere via symbiotic fixation of N_2_ and other gaseous sources through stomata) and the discrimination during uptake, e.g. by mycorrhizal symbioses (Hobbie & Colpaert, 2003; Craine *et al*., 2015).

European silver fir (*Abies alba* Mill.) is one of the most heavily browsed species of the commercially important trees in European forests. Its browsing damage is clearly higher compared to species, such as *Picea abies* and *Fagus sylvatica* (Gill, 1992; Senn & Suter, 2003). Due to its deeper rooting system, silver fir is likely superior to the latter two species to cope with drought stress (Dyderski *et al*., 2018; Vitasse *et al*., 2019; Frank *et al*., 2015; Tinner *et al*., 2013), even though some authors debate its resistance to drought (Battipaglia *et al*., 2009; Vitali *et al*., 2017; George *et al*., 2015). How silver fir can cope with changing climatic conditions may depend on the local climate and soil conditions. Several studies reported high adaptive potential and divergence among its populations in growth, phenology and morphological traits (Hansen & Larsen, 2004; Vitasse *et al*., 2009; Kerr *et al*., 2015; Frank *et al*., 2017). In addition, adult needle *δ*^13^C appeared to be a good indicator of growth rate and timing of seedlings in a common garden (Csilléry *et al*., 2020b).

In this study, we assessed the physiological response of silver fir seedlings to simulated ungulate browsing two growing seasons after the treatment in an ongoing common garden experiment (Frank *et al*., 2017; Kupferschmid & Heiri, 2019; Csilléry *et al*., 2020b). In parallel, we assessed the same physiological traits in the source populations, whenever possible on the mother trees of the seedlings, in their home environments. Our aims were (i) gaining a deeper understanding of the physiological response to browsing stress and (ii) detecting spatially varying selection to physiological traits that may indicate seedlings’ capacity of recovery from browsing stress. Previous analysis of growth traits in the same common garden experiment showed that seedlings recovered from the stress caused by simulated ungulate browsing when only their terminal buds were browsed, but not after the removal of several shoots (i.e. heavy browsing) Kupferschmid & Heiri (2019). Here, we hypothesized that the effect of simulated browsing on physiological traits related to C and N balance would be largely diminished. Further, we expected that physiological traits related to C balance would be correlated with growth traits and would be heritable, while traits related to the N balance would have a weaker genetic component and determined to a larger extent by the local environment. We also tested the specific hypothesis that there is a growth–storage trade–off and if it could be triggered by browsing stress. Previous analysis of growth traits in the same common garden experiment also showed that there is evidence for spatially varying selection pressure for growth and phenology traits (Csilléry *et al*., 2020b). We hypothesized that physiological traits related to the C balance would also be under spatially varying selection, and would be affected by a similar set of environmental variables as growth. In contrast, we expected that traits related to the N balance would be largely environmentally determined. Finally, we hypothesized that physiological traits in seedlings in a common garden mirror the physiology of adult trees in-situ at comparable traits.

## Materials and Methods

### Experimental design and sampling

Our study builds on a large scale common garden experiment aimed at testing growth and phenological differences, and their potential climatic drivers, among Swiss provenances of three major tree species, including silver fir (*Abies alba* Mill.) (Frank *et al*., 2017). The experiment started in 2010, when seeds from three dominant trees per growing site (subsequently called population) were sown in nursery beds at the premises of the Swiss Federal Research Institute WSL (Birmensdorf, Switzerland). In 2012, 16 randomly selected seedlings per mother tree were planted in the open field site near Matzendorf (Swiss Jura Mountains) in a random block design (Fig. 1A and B). In March 2015, at the start of the third field growing season, a simulated browsing experiment started to test seedlings’ morphological response and capacity of recovery to ungulate browsing (Kupferschmid & Heiri, 2019) (Fig. 1B and C). Blocks were randomly assigned to three different treatments: the uppermost buds of the leader shoot were clipped in six blocks (Terminal bud removal), the whole leader shoot and part of the side shoots were clipped in five blocks (Heavy browsing), while the remaining five blocks were left as Controls (Kupferschmid & Heiri, 2019) (Fig. 1B and D; note that Kupferschmid & Heiri (2019) called Terminal bud removal “Light browsing”).

**Figure 1:**
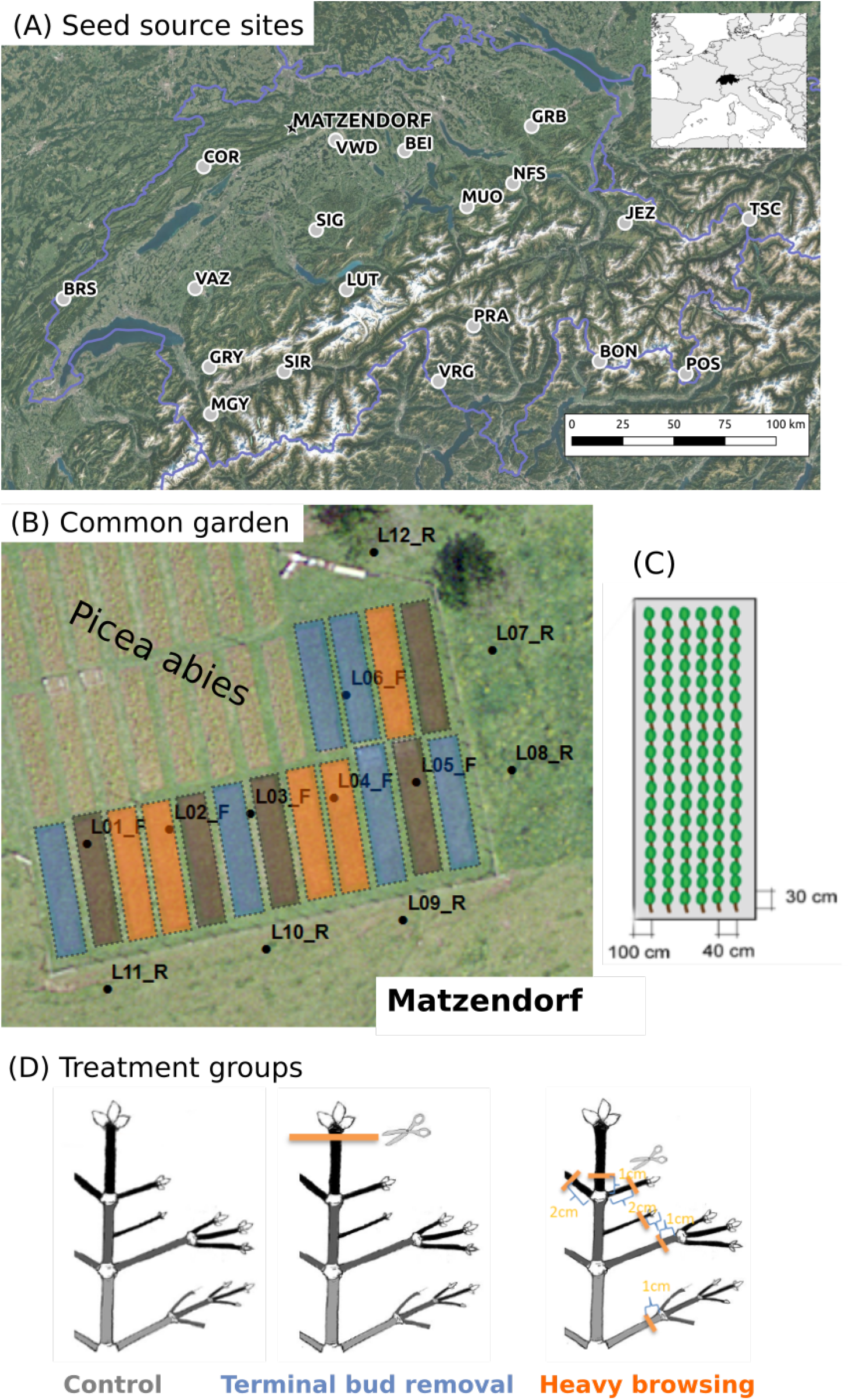
(A) Situation of the 19 seed source populations and the common garden in Matzendorf. (B) Areal photo of the common garden site. Silver fir blocks are highlighted in colors according to the treatment applied (Control - grey, Terminal bud removal - blue, Heavy clipping - orange). Black points labelled starting with “L” indicate the location of the soil samples within and outside of the experimental area. (C) Arrangement of seedlings in a single block. (D) Graphical illustration of the different treatments.

In this study, we measured physiological traits on 19 silver fir populations out of the 90, present in the above common garden study (Table S1, Fig. 1A). Populations were selected to represent the main climatic regions of Switzerland, and were identical to those studied in Csilléry *et al*. (2020a). Needle traits were measured both on the seedlings growing in the common garden and on adult trees of the seed source populations. In September 2016, we sampled seedlings for 2016-grown, approximately 2 cm long, lateral shoots. Adult tree populations were revisited in April 2016 to collect needles from ten trees per site, including the three mother trees, if they could be identified, and other dominant trees from the stand (Csilléry *et al*., 2020b). Approximately, 200 m distance was kept between sampled trees to assure that the environmental heterogeneity of the site was captured, and between population variability reflect real differences as opposed to local growing conditions of individual trees. 2015-grown, light exposed, abaxial needles were selected to assure homogeneity of sampling among trees (Brendel *et al*., 2003). Needles were stored in plastic bags at 5°C and were lyophilized for at least 48 hrs within 24 hrs of their collection.

### Growth traits in seedlings

Growth traits were measured yearly from 2012 to 2016 after growth cessation, and have been analyzed across 90 provenances in Frank *et al*. (2017) and Kupferschmid & Heiri (2019). We re-analysed the 2015 and 2016 growth traits here from the selected 19 populations (i) to check if the effect of browsing using 19 vs 90 populations agrees, and (ii) to assess the relationship between growth and physiological traits. Height was measured from the ground to the highest point of the tree (Height; all trait names are capitalized) or to the tip of the terminal shoot (Terminal Height). Diameter was measured 2 cm above the soil surface. In February 2017, all seedlings were cut 2 cm above the soil surface and their Fresh Weight was determined using a hanging scale (Kern HDBH 5K5N) with a precision of 5 g. Additionally, the weight of 1000 seeds from each mother tree and their diameter at breast height (DBH) were measured to account for potential maternal effects (see details in Csilléry *et al*. (2020b)).

### Sugar, Starch and NSC in seedlings

We measured Sugar in the harvested needles, i.e. the amount of low molecular weight sugars (glucose, fructose and sucrose) by converting them to glucose following the protocol of Wong (1990) and Hoch *et al*. (2002). 8-10 mg of dried ground needles were boiled in 2 ml distilled water for 30 minutes. An aliquot of 200 µl was treated with Invertase and Isomerase from baker’s yeast (Sigma-Aldrich, St. Louis, MO, USA) to degrade sucrose and convert fructose into glucose. The total amount of glucose was determined photometrically at 340 nm in a 96-well microplate photometer (HR 7000, Hamilton, Reno, NE, USA) after enzymatic conversion to gluconate-6-phosphate (hexokinase reaction, hexokinase from Sigma Diagnostics, St. Louis, MO, USA). NSC (Non Structural Carbohydrates) is the sum of low molecular weight sugars and starch. We measured NSC by digesting all starch into glucose, and determined the amount of glucose photometrically. To digest the starch, we used a 500 µl aliquot of the boiled material and incubated with a fungal amyloglucosidase from *Aspergillus niger* (Sigma-Aldrich, St. Louis, MO, USA) for 15 h at 49°C. Starch was derived as the difference between NSC and Sugar. Pure starch and glucose, fructose and sucrose solutions were used as standards, and a standard plant powder (Orchard leaves, Leco, St. Joseph, MI, USA) as a control. NSC concentrations are expressed on a percent of dry matter basis. All samples were analyzed in the same laboratory and by the same person at the Swiss Federal Institute WSL using the same protocol for processing samples in order to minimize biases (Quentin *et al*., 2015).

### Stable isotope traits in adult trees and seedlings

*δ*^13^C, *δ*^15^N, and C and N concentration were measured in the lyophylized needle tissue following the same protocol in adult trees and seedlings. Approximately 80 mg of lyophilized needle material was milled in 2 ml polypropylene tubes equipped with a glass ball (diameter of 5 mm) for 4 min at 30 Hz). Milled samples were directly weighted in small tin capsules (approx. 5 mg, XPR2 microbalance from Mettler Toledo), and combusted in an elemental analyzer (Flash EA by Thermo Finnigan, Bremen, Germany) coupled to an isotope ratio mass spectrometer (Delta XP by Thermo Finnigan, Bremen, Germany) by a Conflo II interface (Thermo Finnigan, Bremen, Germany). All C and N isotope values are reported using the Vienna-Pee Dee Belemnite (V-PDB) standard using the *δ*-notation following Werner & Brand (2001).

### Environmental data

We characterized the environmental conditions at the home sites of the 19 populations and the soil of the common garden site. First, a soil profile was taken at each of the 19 seed source sites and soil N and C concentrations [%] were determined from the uppermost part of the A horizon (see Frank *et al*. (2017) for more details). We also placed soil profiles at several locations at the common garden site (see Fig. 1B) and determined the C and N concentrations [%] across different depth. Second, we determined the latitude, longitude, elevation, slope, topographic wetness index (TWI) and aspect of each site from a 100m resolution digital terrain model. Third, we extracted raw climate time series from CHELSAcruts (http://chelsa-climate.org/chelsacruts/), which is a time series version of the CHELSA data (Karger *et al*., 2017). Several climatic indices were calculated across the 1901-1979 period, thus excluding the recent years that are affected by climate warming and did not affect the establishment of current adult trees.

### Statistical analysis

Seedlings that died during the experiment or were damaged by frost or insect herbivory were omitted (N=15). We also excluded outlier observations that were not consistent with the treatments as well as suggested observation errors. In particular, we excluded control seedlings that had a height loss between 2014 autumn and 2015 spring (N=3), seedlings with Terminal Bud removal that had a height loss greater than 20% (N=7), and seedlings in the Heavy Browsing treatment that exhibited no height loss due to the treatment (N=2), leading to a total of 224 seedlings in the Control, 271 in the Terminal bud removal, 218 in the Heavy browsing treatment. The concentration of Starch was calculated based on NSC minus free sugars, which led to some negative values. For the sake of easier interpretation of the effects, we added ten to all Starch values to assure that all observations are non-negative. Starch, Height, Terminal Height and Fresh Weight were log-transformed to achieve a close to normal distribution. All other traits were normally distributed based on visual evaluation of histograms. We also calculated a derived trait from the different height measures to check the homogeneity of the treatments on the targeted seedlings:

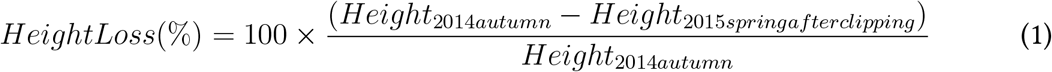

First, we used a linear mixed-effects model, so-called animal model (Henderson, 1975), implemented in the R package ASReml-R that uses ASReml version 4.0 (Butler *et al*., 2009) to estimate the proportion of the trait variance explained by the treatment, block, population of origin, and genetic (family) effects. We fitted a separate model to Control and Terminal bud removal together and Control and Heavy browsing together for two reasons. First, a treatment variance component would have been difficult to interpret with the two treatments together in one model. Second, we wanted to quantify the effect of Terminal bud removal and Heavy browsing separately on the growth and physiology of the seedlings. We used the following model:

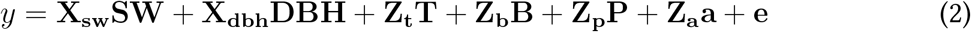

where y is a vector of observations for a trait on all seedlings, and **X** and **Z** are incidence matrices relating the covariates and random effects to the observations, respectively. **SW** and **DBH** are covariates for the maternal effects Seed Weight and Diameter at Breast Height, respectively. The random effects were **t** for treatment (Terminal bud removal or Heavy browsing), **b** for block, **p** for populations, while **a** is a vector of individual breeding values with variance *V ar*(*a*) = *A*×*V*_*A*_, where *A* is the inverse kinship matrix constructed based on half-sib family relationships using the function *ainverse*, and *V*_*A*_ is the additive genetic variance, i.e. part of trait variance that is due to heritable genetic factors. Finally, **e** is the vector of residuals following *E* ∼ *N* (0, *V*_*E*_), where *V*_*E*_ is the error variance. In order to evaluate the significance of the covariates, we compared models with and without these using a Wald-test (wald.asreml function). We also compared models with and without random effects using a likelihood ratio test (p-values based on the *χ*^2^ distribution are reported). All variance components, including treatment, block, population and genetic, were expressed as proportions to the total phenotypic variance, *V*_*T*_. The proportion of the trait variance due to genetic factors is the heritability of the trait, denoted as *h*^2^ = *V*_*A*_*/V*_*T*_ (Falconer & Mackay, 1996). The significance of variance components was assessed using z-scores with z>2 indicating a non-zero variance component.

Second, the model including the Control and Terminal bud removal groups revealed that including the treatment did not improve the model fit (with the exception of Terminal Height in 2015 and 2016, and C concentration), and even when it did, treatment did not explain a significant part of the trait variance (with the exception of Terminal Height in 2015). Thereby, we applied a simpler model excluding treatment effect for a data set pooling the Control and Terminal bud removal groups.

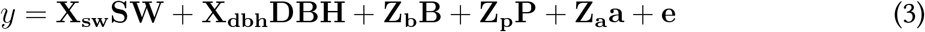

We used this model to estimate the amount of population differentiation that is due to genetic factors, defined as *Q*_*ST*_ = *V*_*P*_ */*(*V*_*P*_ + 2*V*_*A*_), where *V*_*P*_ is the population variance (Whitlock, 2008). Estimating *Q*_*ST*_ requires a large sample size, so we benefited from being able to pool together the two treatment groups to obtain more reliable estimates. Previous analyses of this common garden experiment have already estimated *Q*_*ST*_ for several growth and phenology traits from 2013 and 2014 (Frank *et al*., 2017; Csilléry *et al*., 2020b). We took advantage of having the full time series of growth traits from 2012 to 2016 to assess the evolution of *Q*_*ST*_ in time, and to contrast *Q*_*ST*_ between growth and physiological traits. Additionally, we used the genetic marker data available from Csilléry *et al*. (2020b) to test if trait divergence between populations was significantly higher than that at genetic markers (*F*_*ST*_) using the R package *QstFstComp* (Gilbert & Whitlock, 2015).

Third, we assessed the role of environment in driving trait divergence between populations in seedlings. We only used traits that expressed a *Q*_*ST*_ significantly higher than zero, i.e. Height, Starch and *δ*^15^N. Although the effect of environmental variables on Height have been assessed by earlier studies (Frank *et al*., 2017; Kupferschmid & Heiri, 2019; Csilléry *et al*., 2020b), we repeated these analyses for the sake of completeness and also to see if findings of earlier studies were confirmed despite the reduced sample size. For these tests, we extracted the population effects from the pooled model (i.e. Control + Terminal bud removal) and correlated these with the environmental variables using a Spearman correlation test (see details below). Additionally, we explicitly tested if there was a growth–storage trade–off by correlating the Height in 2015 with the difference in Starch between Control and Heavy Browsing treatment (population means). Finally, we also tested the correlation between traits measured in adult trees *in-situ* and environmental variables. In order to do so, we first tested if traits were significantly different between populations, and since they were (Kruskal-Wallis tests trait-by-trait: *χ*^2^ > 44.86, df=18, p-value<0.001, see Fig. S2 for details), we could use all five traits measured in adult trees, i.e. C concentration, *δ*^13^C, N concentration, *δ*^15^N, and C/N for this analysis. Note that the same test was already performed for *δ*^13^C by Csilléry *et al*. (2020b).

For all traits measured in seedlings or adult trees, we tested their correlation with a total of 37 environmental variables including topographic and bio-climatic variables, drought and frost indices, and soil variables extracted from local soil pits (Table S5 and S6). We used a correction for multiple testing that accounts for the correlation between variables, thereby for the non-independence of tests. A Principal Component (PC) analysis of all environmental variables (*prcomp* function in R using scale=TRUE) revealed that nine PC axes explain 95% of the total variance (94.75%), thus we adjusted the p-values using a Bonferroni correction as if we performed nine independent tests for each trait. Further, we used an even more strict correction: we accounted for testing seven traits (in seedlings) and five traits (in adult trees), which could be considered as two and four independent tests based on the same argument as above (i.e. two axes explained 99.9% and four axes 99.64% of the total variance in seedlings and adult trees, respectively). Thereby we adjusted for 18 and 36 independent tests in seedlings and adult trees, respectively.

## Results

### Response to simulated browsing

Terminal bud removal affected the growth of seedlings one growing season after the clipping experiment (i.e. in 2015 autumn): their Terminal Height was reduced, but not their Height or Diameter (Table S2, Fig. S1 and 2). In contrast, Terminal bud removal did not explain a significant part of the variation neither in growth traits nor in physiological traits two growing seasons after the clipping experiment (i.e. in 2016 autumn) (Fig. 2), although the model fit still improved by including the Terminal bud removal treatment for Terminal height and C concentration (Table S2). In contrast, heavy browsing had a long-lasting effect on most traits: both Height and Terminal Height decreased (Fig. S1), and both in 2015 and 2016 over 40% of the trait variation was explained by the treatment (Fig. 2). In contrast, Heavy browsing did not explain trait variation in 2015 Diameter and 2016 Diameter and Fresh Weight (Fig. 2). Among the physiological traits, only *δ*^13^C and *δ*^15^N, were affected by the Heavy browsing treatment: over 40% of the trait variation was explained by browsing stress even though the standard error of this variance component was high (Fig. 2 and 3). On average, *δ*^13^C decreased by 0.86 ‰ as a result of the Heavy browsing treatment, with the greatest decrease observed in population POS, with 1.39 ‰ (before last population on Fig. 3). *δ*^15^N increased, on average, by 1.07 ‰ as a consequence of Heavy browsing, with an increase being as high as 1.79 ‰ in SIG (fifth population on Fig. 3).

**Figure 2:**
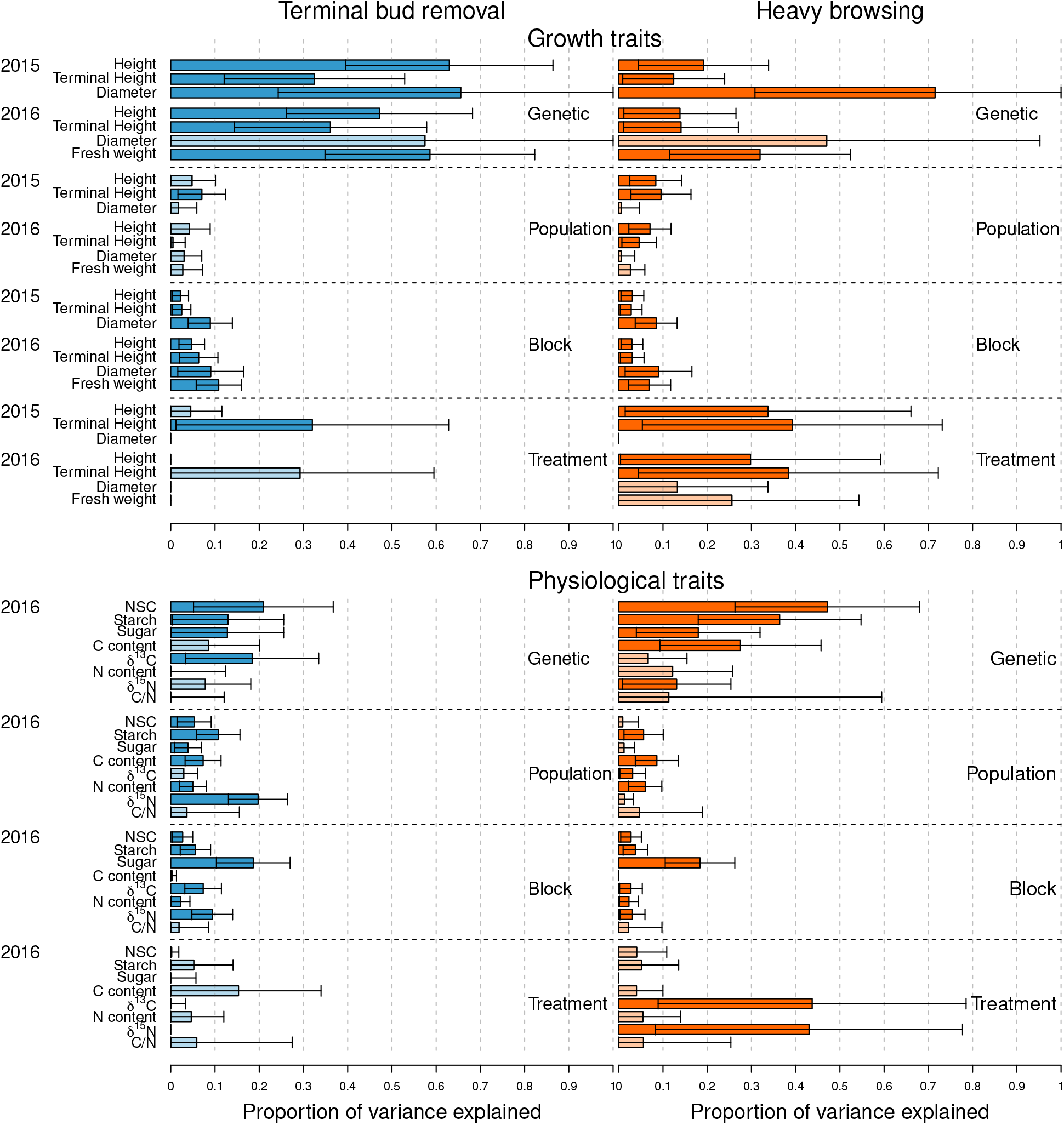
Variance components from the mixed effects models described in equation 2 expressed as proportions of the total variance. The first column in blue shows the model including the Control and the Terminal bud removal groups, and Treatment shows the trait variance explained due to Terminal bud removal. The second column in orange shows the model including the Control and the Heavy browsing groups, and Treatment shows the trait variance explained due to Heavy browsing. Dark color bars indicate that a significant part of the trait variance was explained by the given factor (Genetic etc.), while light color bars indicate non-significant variance components.

**Figure 3:**
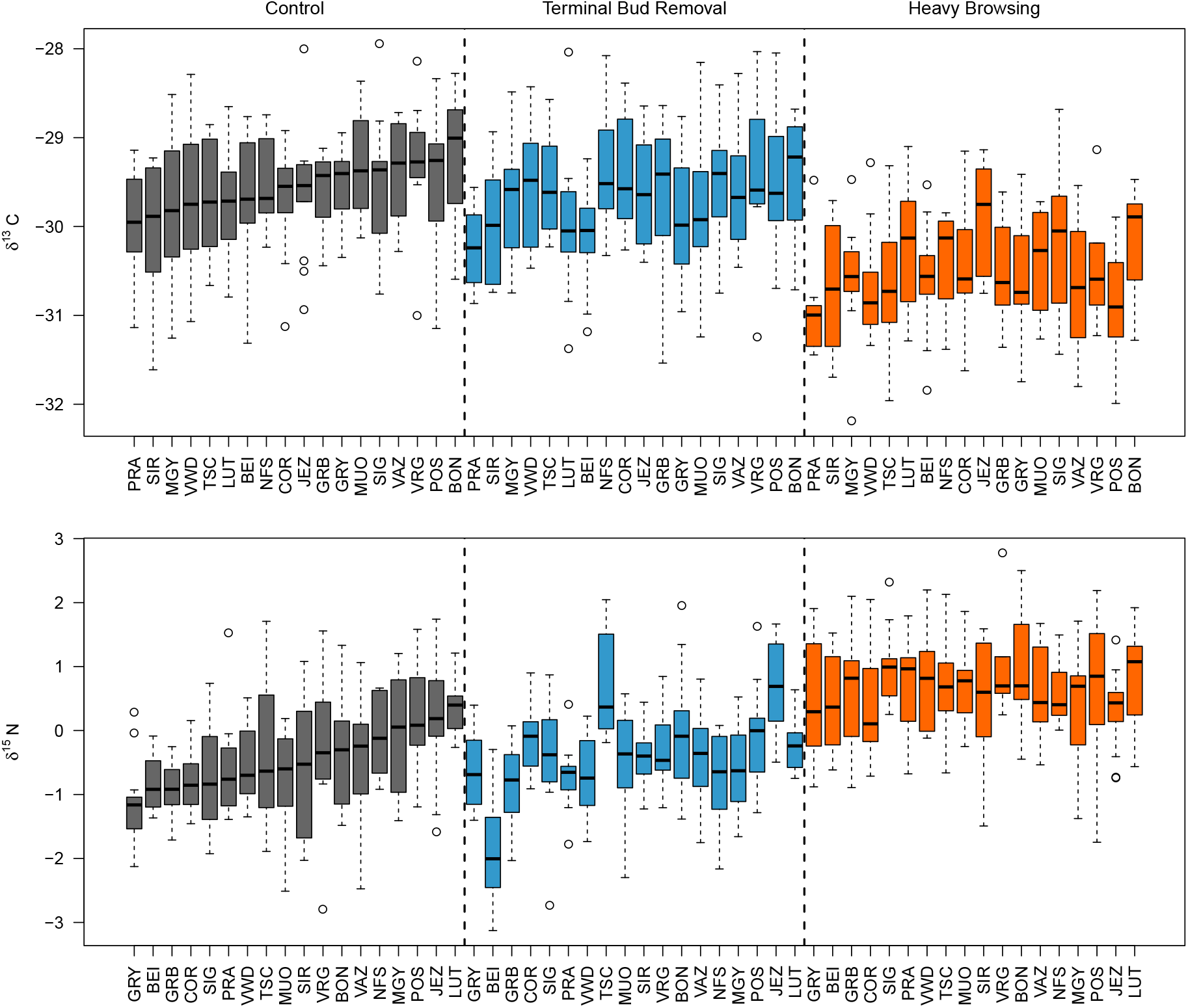
Boxplots of the raw *δ*^13^C and *δ*^15^N values per population across the three treatment groups: Control, Terminal bud removal, and Heavy browsing (see Fig. 1). Populations are ordered according to their medians in the Control group.

Local environmental variation may have interfered with the treatment within the common garden, and was quantified using Block effects. Including Block significantly improved the model fit for all traits but C concentration (Table S2). Block explained the highest proportion of trait variation in Fresh weight (10.8%, SE: 5.0%) among the growth traits, and in Sugar (18.6%, SE: 7.6%) among the physiological traits. Block variance was, on average, higher than Population variance for growth traits (Fig. 2), suggesting that the local growing conditions were just as important as the population of origin.

### Genetic and population effects on trait variation

Genetic effects explained the most variation in both growth and most physiological traits, and with or without Heavy browsing stress (Fig. 2, Table S3). The heritability of growth traits varied between 0.65 (Diameter 2015) and 0.38 (Terminal Height 2015). The genetic variance component was also significant for Diameter in 2016 (*h*^2^=0.47, SE: 0.21) when the Terminal bud removal treatment was removed from the model (unlike in the model shown on Fig. 2). Among the physiological traits, the highest heritability was observed for NSC (*h*^2^=0.21, SE: 0.16) and *δ*^13^C (*h*^2^=0.18, SE: 0.15). Interestingly, the genetic effects became stronger under the Heavy browsing treatment, especially for storage related traits, thus NSC, Starch, Sugar and C concentration (Fig. 2, Table S3). In contrast, the genetic effects lessened under stress for *δ*^13^C (Fig. 2). Among the two proxies for maternal effects, Seed weight had a significant effect on all growth traits and a marginally significant effect on C concentration (Table S2). In contrast, the size of the mother tree (DBH) did not affect seedling performance in any way (Table S2).

Population of origin explained a significant part of the trait variance only for Terminal height among the growth traits, however, under Heavy browsing stress, the population differences became stronger and significant for all growth traits, except for Diameter and Fresh weight (Fig. 2). Taken together the high genetic and low population effects (without browsing stress, using equation 3), *Q*_*ST*_ s were not significantly different from a neutral expectation derived from genetic markers for the 2015 and 2016 growth traits (see Fig. 4 for Height, and Table S4 for other the traits). Note that higher *Q*_*ST*_ values were detected for Height in previous years by previous studies: *Q*_*ST*_ was significantly different from zero for 2013 Height using 90 populations in Frank *et al*. (2017) and *Q*_*ST*_ was significantly larger than *F*_*ST*_ for 2013 and 2014 Height using 19 populations in Csilléry *et al*. (2020b). These trends were also confirmed herein despite the reduced sample size due to mortality and the browsing experiment (Fig. 4 and Table S4).

**Figure 4:**
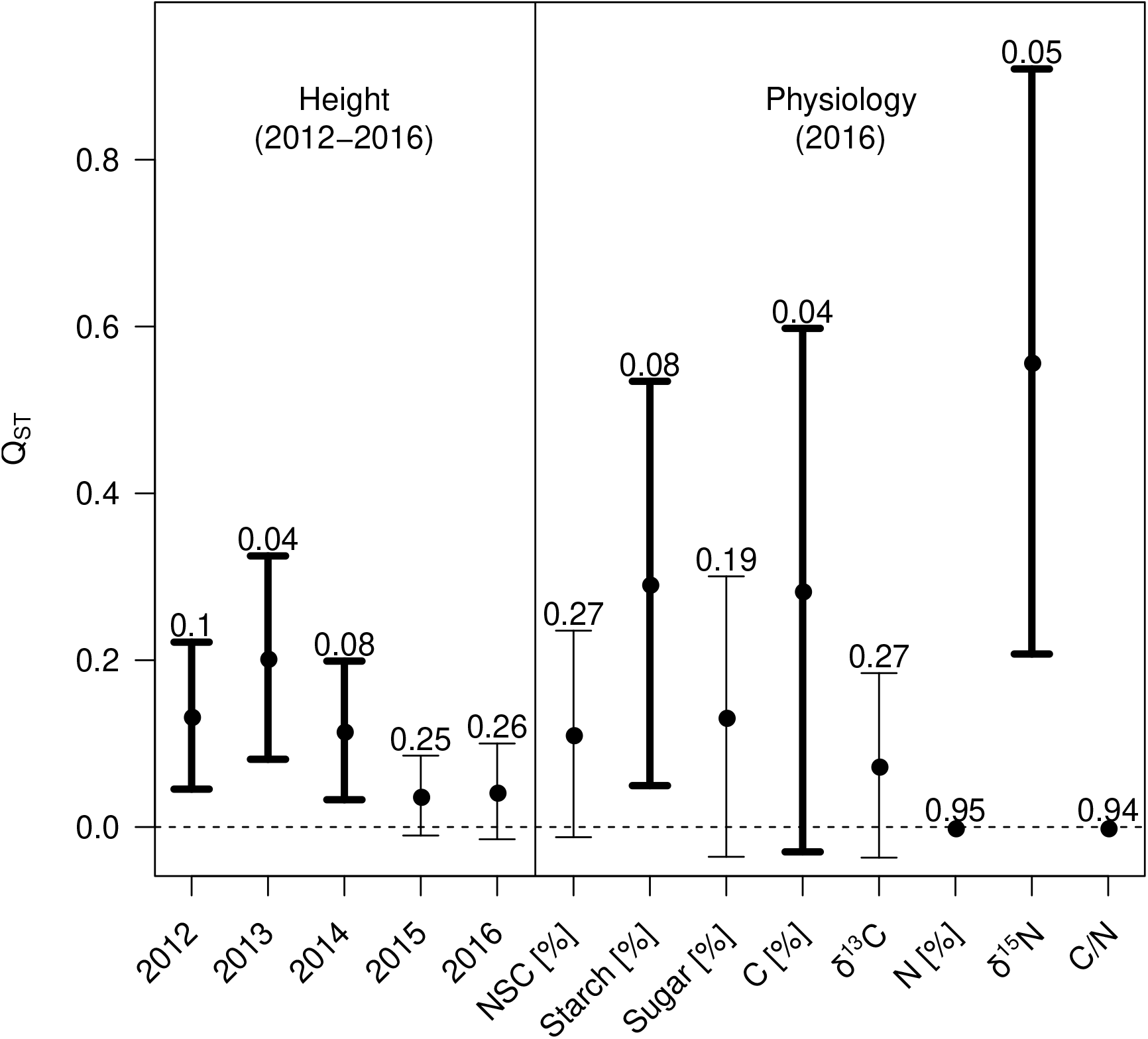
Population genetic differentiation (*Q*_*ST*_) estimated from the pooled model (equation 3), combining data from the Control and Terminal bud removal groups for Height from 2012 to 2016, and for physiological traits measured in 2016. The numbers above each bar show the p-values from the *Q*_*ST*_ -*F*_*ST*_ test (see full test results in Table S4). Traits that showed evidence for spatially varying selection shown in bold.

A higher proportion of the trait variation was due to population of origin in physiological than in growth traits (Fig. 2). Further, all population effects for physiological traits were significant except C concentration and C/N (Table S2 and Fig. 2). *δ*^15^N had an exceptionally high proportion of trait variance explained by population (19.7%, SE: 6.7%). Some of these population differences might have been the result of spatially varying selection at the source populations. Indeed, the *Q*_*ST*_ of Starch and *δ*^15^N were significantly different from zero (Fig. 4). When comparing *Q*_*ST*_ to a neutral expectation derived from genetic markers based on *F*_*ST*_, we found evidence for selection on Height 2013 and C concentration, and tendencies for Height on other years, *δ*^15^N and Starch (Table S4). However, note that the mixed effects model used by *QstFstComp* did not include Block and covariates for maternal effects, which may have altered the results.

### Environmental drivers of population divergence and response to browsing

Several environmental variables related to the temperature of the home sites were correlated with seedlings’ growth in the common garden (Table S5). Generally speaking, warmer and more thermally stable home sites were related to faster growth, as it has been shown by previous analyses of data from this experiment (Frank *et al*., 2017; Kupferschmid & Heiri, 2019; Csilléry *et al*., 2020b). More interestingly, we found evidence for a growth–storage trade–off at the level of populations mediated by browsing stress: seedlings from fast growing populations, which often came from warm places, tended to decrease their storage (Starch) in response to heavy browsing stress, while seedlings from slow growing populations, often originating from cold places, tended to increase their Starch levels when heavily browsed (Fig. 5). Note that the correlation was also significant for family means (Pearson correlation, r=0.33, p-value=0.018), suggesting a potential genetic underpinning for this trade–off. Finally, the population effects for Starch under non-stressed conditions (model equation 3) did not reveal correlations with any of the environmental variables (Table S5), suggesting that the growth–storage trade–off is triggered only under heavy browsing stress.

**Figure 5:**
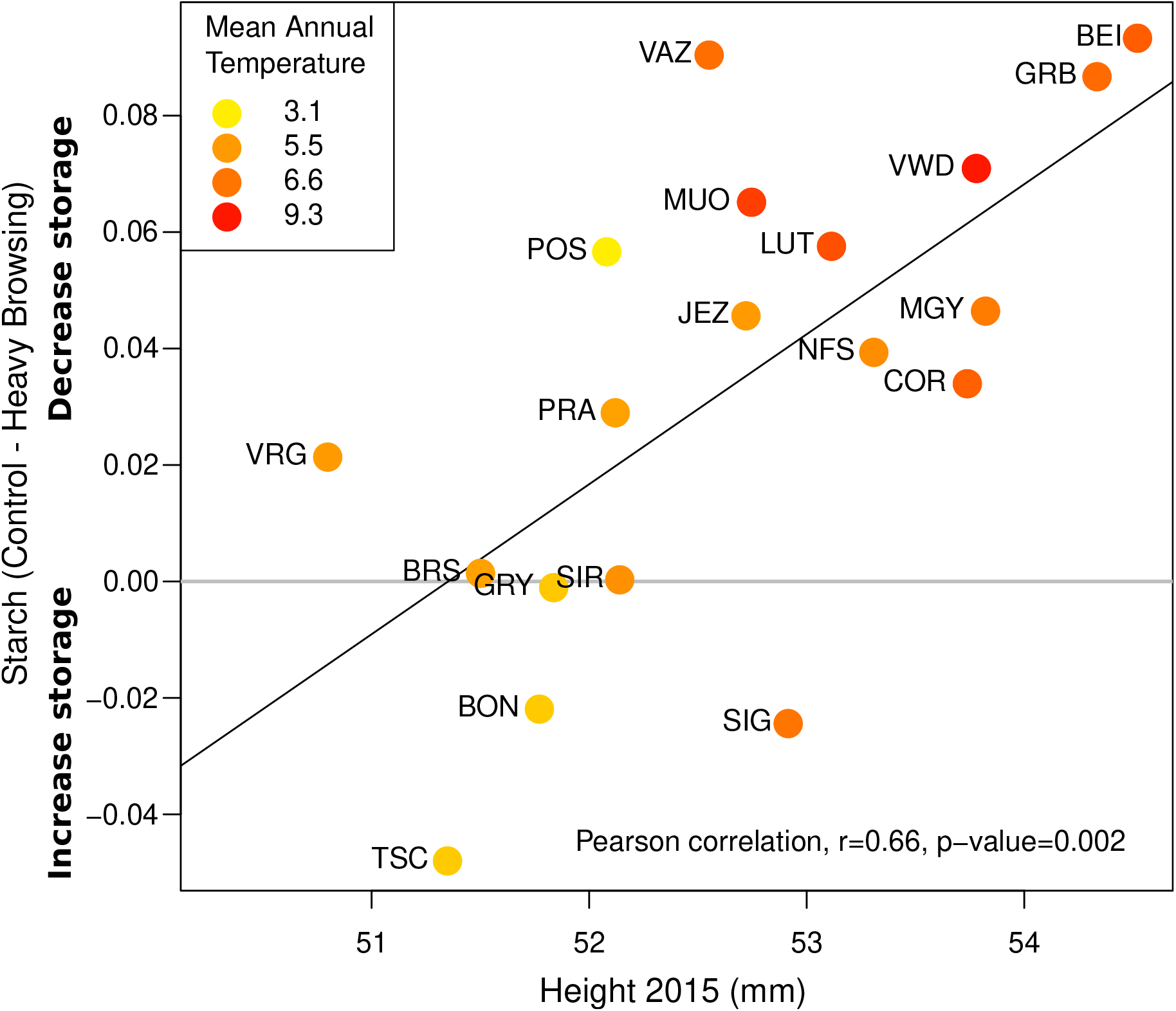
Growth–storage trade–off according to climate origin. Each point shows the difference between population means for Control and Heavy browsing treatments against the mean Height of the population in 2015, after the browsing treatment. Color code corresponds to the mean annual temperature of the home sites. See Table S5 for correlation between Height and climatic variables.

The common garden site was within the species distribution range, and was climatically close to many home environments (Frank *et al*., 2017). However, using soil samples from the common garden sites, here we showed that the soil N concentration was higher and the C/N (mean=10.6) was lower in Matzendorf than in any of the 19 home sites (mean C/N of 16.4, Fig. S3). Further, we found that forest soils *in-situ* had a large variation across Switzerland (Fig. S3). The most N poor soils were observed in mountain populations (top four sites: PRA, TSC, BON, SIR), and the most N rich soils at the Swiss Plateau (top four sites: VAZ, COR, BEI, GRB) (Fig. S3 and Fig. 6). The greater between-site environmental variation *in-situ* was reflected by a higher coefficient of variation (CV) in the population medians of traits measured in adult trees in comparison to the CV of the seedling population effects (Fig. 6 and Fig. S4). Interestingly, heavy browsing stress also increased the variation across populations: the CV for C/N and *δ*^15^N was almost as high for heavily browsed seedlings as for adult trees *in-situ* (Fig. 6).

**Figure 6:**
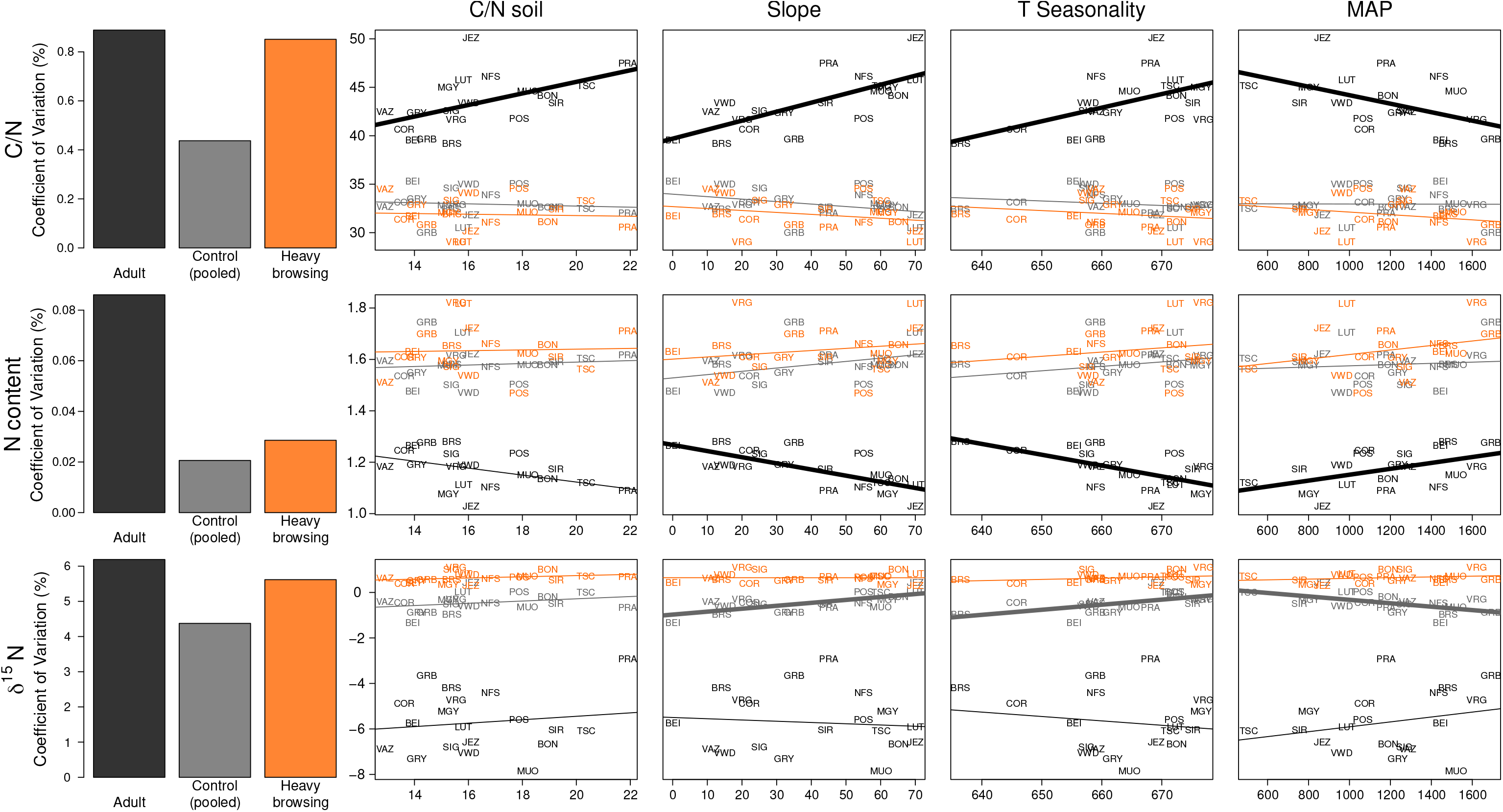
The strongest significant correlations between environmental variables (x axes) and mean trait values (y axes) in seedlings (orange and grey lines) and in adult trees (black lines). Barplots in the first column show the coefficient of variation among populations. Slope of the terrain is expressed as a percentage. T stands for temperature and measured in °C. MAP stands for Mean annual precipitation (in mm). All correlation tests between environmental variables and trait values with correction for multiple testing are shown in Tables S5 and S6. Thick lines indicate significant correlations while thin lines indicate non-significant relations. For simplicity, MAP is shown instead of other precipitation variables that had more significant correlations with trait values. Note that for seedlings, correlation tests were performed on population effects extracted from mixed-effects models, while this figure shows simple population means for an easier comparison with adult values.

Although soil composition may vary rapidly within short distances, we found that the analysis of samples from a single soil profile *in-situ* explained the variation in traits measured in adult trees. Soil C/N was strongly and positively correlated with the population median C/N from adult tree needles (Fig. 6, Table S6). Population median C/N and N in adult tree needles was significantly associated with the Slope (and the Topographic wetness index), such that steeper terrain led to lower N concentration, thus higher C/N, in needles. Additionally, higher temperature stability (T seasonality) and higher annual precipitation (MAP) were also associated with higher N concentration (and lower C/N) (Fig. 6, Table S6). Further, seedling’s population effects for *δ*^15^N were correlated with principally the same environmental variables as N concentration and C/N in adult trees (Tables S5 and S6), such as with the Slope, the Topographic wetness index, temperature seasonality and precipitation variables (Fig. 6, Table S5). Thus, it appeared that descendants of populations that came from steep terrains, thus less developed soils, did not discriminate the two N isotopes in the common garden (Fig. 6).

## Discussion

### Heritability and browsing stress in growth and physiological traits

Multi–site experiments or experiments involving different treatments allow to detect genotype–environment or genotype–treatment interactions, but often lead to reduced heritability estimated or significant heritability in one site or under one treatment only (Grattapaglia *et al*., 2018). Here, we performed a simulated browsing experiment with two treatment levels, which allowed us to explore how the genetic component of trait variation is altered by stress. Genetic factors explained most variation in growth traits (on average, 51.5%), but only 10.2%, on average, in physiological traits under control conditions (Fig. 2, Table S3). In contrast, heavy browsing stress considerably reduced the genetic effects on growth (to 30%, on average), but doubled those on physiological traits related to storage (Fig. 2).

In this study, the relatively low heritability in physiological traits could be attributed to factors that reduce the additive genetic variance or those that increase the environmental variance. First, some physiological traits might be more closely related to fitness, thus natural selection might have removed much of the additive genetic variance (Merilä & Sheldon, 2000; Hansen *et al*., 2011; Hoffmann *et al*., 2016). It is difficult to find general evidence for such a phenomenon, but, for example, previous works that studied traits closely related to fitness, such as seed production on oaks, found a heritability higher than that for growth traits (Caignard *et al*., 2019). Second, some physiological traits are less integrative than growth traits, and they change on shorter time scales (Millard & Grelet, 2010). Indeed, the most integrative physiological traits had the largest part of the trait variation explained by genetic effects, such as NSC and Starch and the *δ*^13^C (Fig. 2). Finally, physiological traits appeared to be more effected by micro-environmental variation, as suggested by the relatively high block effects, for example for Sugar (Fig. 2).

Several previous common garden studies estimated the heritability of growth traits and found moderate to high heritabilities (Cornelius, 1994), even though, it is well-known that these values are inflated and likely much lower in natural settings (*e*.*g*. Latreille & Pichot, 2017). Fewer studies estimated the heritability of physiological traits, and most of them concentrated on *δ*^13^C or water use efficiency (WUE), and usually from wood, and not from needles. For example, in Maritime pine, Brendel *et al*. (2002b) found a moderate heritability for *δ*^13^C based on a bulk sample across several tree rings (0.17), and Marguerit *et al*. (2014) found a higher heritability for *δ*^13^C (0.29), compared to circumference and height. Here, we found a heritability of 0.18 (SE: 0.15) for *δ*^13^C under control conditions (Table S3), but the genetic effects vanished for seedlings under browsing stress (Fig. 2).

Traits related to the N balance are often considered principally environmentally determined, and, as a result, very few studies assessed the genetic factors that may influence them. Some previous studies assessed the so-called nitrogen use efficiency (NUE) in fertilization experiments, defined as added N per stem biomass (Li *et al*., 1991) or approximated using C/N and variation in *δ*^15^N (Hu *et al*., 2021). Li *et al*. (1991) found that NUE traits were under a moderate to high genetic control in Loblolly pine: families with higher NUE had greater root length and stem height at low N concentrations, but not at high N concentrations. Xu *et al*. (2003) evaluated the heritability of *δ*^15^N in hoop pine; the first study assessing the genetics of this trait in forest trees. They found that under water stress, *δ*^15^N was higher and had a moderate heritability, but there were no significant genetic effects at the wet site. The same trends were confirmed for European beech in a watering experiment, i.e. higher *δ*^15^N (and also lower *δ*^13^C) under water stress, and a family treatment interaction for traits related to N concentration (Aranda *et al*., 2017). Finally, most recently, Hu *et al*. (2021) found genetic and clinal environmental variation for N isotope discrimination in heart-leaved willow. In agreement with these studies, here we found a significant genetic variation for *δ*^15^N only under browsing stress, which has similar physiological effects to drought stress (see below). Our results also confirm that under stress seedlings discriminate N less (higher *δ*^15^N, Fig. 3). These results suggest that genetic studies of *δ*^15^N could enhance our understanding of N acquisition and metabolism in forest trees (Hu *et al*., 2021). Finally, trees may also influence the composition of microbial communities, thereby the microbial biomass N beneath them. Schweitzer *et al*. (2008) found that individual genotypes in *Populus* explained up to 70% of the variation soil microbial community composition. Thus, part of the genetic variance for N related traits might reflect a dynamic interaction between the trees and soil microbial communities.

### Response to simulated browsing

Loosing the Terminal bud, thus the apical meristem tissue, did not have long-term impacts on the growth and carbon and nitrogen storage and re-mobilization in seedlings: two growing seasons after the loss they did not differ from control seedlings in the measures traits, with the exception of Terminal height (Table S2 and Fig. 2). In contrast, Heavy browsing had three key long lasting effects affecting seedlings’ growth and storage, water use efficiency, and N balance. We discuss these effects in the following paragraphs.

First, seedlings had a reduced height growth, but interestingly, seedling’s diameter and fresh weight were not affected, suggesting that the overall growth of the seedlings had recovered two vegetative seasons after the stress. This is because seedlings altered their growth form: they became shorter but grew more side shoots (multi-stemmed growth), which is a typical reaction to browsing in silver fir (Kupferschmid & Heiri, 2019), but also in other tree species (*e*.*g*. Lehtilä *et al*., 2000). Additionally, we found that the fastest growing populations, originating from the warmest regions, decreased their Starch concentration the most as a reaction to heavy browsing stress, suggesting a potential genetic underpinning for a growth–storage trade–off (Fig. 5). This result supports the idea that non-structural carbon (NSC) accumulation occurs at the expense of growth and that there is genetic variation for this trade–off (Palacio *et al*., 2014). Our results are also in agreement with the frequently observed accumulation of NSC in trees under cold and dry conditions, which has been interpreted as a precautionary measure by plants and may be indicative of C limitation for growth (Wiley & Helliker, 2012).

Second, seedlings had a lower water use efficiency (decreased *δ*^13^C) after simulated heavy browsing. The reduced water use efficiency might be due to an increase in stomatal conductance, which would increase photosynthesis and therefore compensate for the loss of leaf surface. A stomatal opening has been shown as a compensatory reaction of plants to a reduction in leaf surface by browsing or leaf detachment (Welker & Menke, 1990). However, the increase in photosynthesis would be smaller than the concurrent increase in transpiration, which would result in the observed decrease in water use efficiency.

Third, simulated heavy browsing increased *δ*^15^N (Fig. 2 and 3). This could be related to the above mentioned increase in transpiration, which could also have increased the transport of nitrate to the leaves, even though we did only observe a slight but not significant increase in needle N concentration as a result of heavy browsing. Even though gymnosperms are known to have low nitrate reductase in the leaves, it has also been shown that it can be induced by providing nitrate (Smirnoff *et al*., 1984), which suggests that nitrogen rich soils (as was the case in our experiment) might increase the nitrate concentration in needles (Smirnoff & Stewart, 1985). As inorganic nitrogen is enriched compared to assimilated, organic nitrogen (Pritchard & Guy, 2005; Cui *et al*., 2020; Hu *et al*., 2021), inorganic nitrate in leaves might perhaps explain the shift to a more enriched total nitrogen isotope composition that we observed for the needles of the heavy browsing treatment.

### C and N isotope discrimination in seedlings vs adult trees *in-situ*

*δ*^13^C in seedlings was not correlated with the population mean *δ*^13^C in adult trees in the home sites after correction for multiple testing (Table S6, Fig. S4). This is probably due to the difference in environmental conditions of the common garden site on the one hand, and the different original sites on the other hand. Strong environmental differences among the original sites are also suggested by the stronger variability of *δ*^13^C among the adult populations, whereas in the homogeneous environment of the plantation site, the range of *δ*^13^C values for the seedlings was lower (Brendel *et al*., 2002a).

Adult needle *δ*^15^N were very low, suggesting low discrimination to ^15^N, which is often the result of a strong mycorrhiza activity in N poor forest soils (Hobbie & Colpaert, 2003; Craine *et al*., 2015). In contrast, in the common garden both soil and needle C/N were much lower than *in-situ*, in agreement with the fact that the common garden was established at an agricultural site that has been fertilized before. Seedling needle *δ*^15^N values were high (zero or positive), suggesting low/no discrimination (Fig. 3). It is likely that seedlings did not develop mycorrhiza, but rather used easily accessible existing N pools, or the fertilizers that have been used were rich in ^15^N. Indeed plants that experience greater N availability may reduce their dependence on mycorrhizal fungi. A reduced dependence on mycorrhiza can enrich plants in ^15^N by reducing the depletion associated with N transfers from mycorrhizal fungi (Högberg *et al*., 2011). Alternatively, we may speculate that due to a reduction in the root to shoot ratio, they were not able to supply sufficient food to their mycorrhiza, which could have also contributed to their increased ^15^N uptake. Indeed, as soil N availability increases relative to C, soil microbial biomass may become more enriched in ^15^N (Dijkstra *et al*., 2008).

### Evidence for spatially varying selection

Population differences for Height, Starch and *δ*^15^N appeared to be the result of natural selection (Fig. 4, Table S4). We confirmed the *Q*_*ST*_ values reported by Frank *et al*. (2017) and Csilléry *et al*. (2020b) in 2013 and 2014 with our reduced data set. However, the population differences diminished with time (Fig. 4). In 2016, in the 8^*th*^ growing season none of the growth traits showed significant population genetic differentiation. This decrease in *Q*_*ST*_ principally stem from a decrease in the within population variance, while the genetic variance component stayed relatively stable over time. This latter is also supported by the fact that our proxy for maternal effects, seed weight, was still significant in 2016 for growth traits (Table S2). In contrast, populations might have become more similar to each other with time because all seedlings were, to some extent, stressed at the common garden site in Matzendorf, as suggested by the restricted range of *δ*^13^C (Fig. S4).

A surprisingly strong correlation was detected between seedling *δ*^15^N and N concentration with the slope of the terrain, and, a slightly weaker but still significant correlation with temperature stability and precipitation (Fig. 6, Table S6 and S5). Seedlings coming from mountain populations growing on steep slopes, great temperature fluctuation and low amounts of precipitation, had a significantly lower N discrimination in the common garden. This finding, along with the significant *Q*_*ST*_ -*F*_*ST*_ tests, indicate that silver populations from mountainous regions across Switzerland have been selected to grow on N poor, potentially less developed and/or drained, soils. This finding was further corroborated by the fact that N concentration in needles of adult trees was lower on steep slopes than on flat ground, strongly indicating that steep slopes are the most N poor environments (Fig. 6). Seed sources from these poor environments generally had a low growth rate and high storage, thus might be slower to recover from browsing stress than fast growing provenances from the warm environments with developed soils, such as the Swiss plateau.

## Conclusions and outlook

Conifers remain dominant only in the most hardy habitats across the globe: they are the most drought and frost resistant trees and can grow on mountain soils poor in nutrients. Thus, it is plausible that conifer populations harbor variation for NUE and related traits, such as *δ*^15^N. Here, we found that N concentration and *δ*^15^N had a strong genetic component and showed evidence for adaptation to spatially varying selection. We encourage future studies on other species and environments to assess these traits as well as soil N concentration and mycorrhizal activity. N acquisition and assimilation play a key role in growth and recovery from drought stress (Li *et al*., 1991; Gessler *et al*., 2017; Millard & Grelet, 2010), thus in recovery from browsing stress.

## Data and Materials Availability

New data published in this study including seven seedling physiological traits (NSC, starch, sugar, C concentration, *δ*^13^C, N concentration, and *δ*^15^N) and three adult stable isotope traits (C and N concentrations, and *δ*^15^N) will be made available in the Dryad Digital Repository upon acceptance. *δ*^13^C from adult tree needles has been published in (Csilléry *et al*., 2020b), and is available in the Dryad Digital Repository.

## Supplementary Data

**Table S1** Names, political and geographic situation of the 19 silver fir populations.

**Table S2** Model comparisons.

**Table S3** Trait heritabilities under control conditions.

**Table S4** *Q*_*ST*_ -*F*_*ST*_ tests.

**Table S5** Correlation between traits measured in seedlings and environmental variables.

**Table S6** Correlation between traits measured in adult trees and environmental variables.

**Figure S1** Height loss due to Terminal bud removal and Heavy browsing treatments per block.

**Figure S2** Physiological trait values in adult trees *in-situ* per populations and Kruskal-Wallis test of population differences.

**Figure S3** C/N ratio of the top mineral soil layer at the common garden site in Matzendorf and at the seed source populations.

**Figure S4** Coefficient of variation in C concentration and *δ*^13^*C*, and correlation between population medians in adult trees (and seedling population effects) and environmental variables.

## Acknowledgements

We thank the ADAPT project for granting us access to the growth data before the simulated browsing experiment. We thank Jens Nitzsche, Barbara Ganser, and Olivier Charlandie for help with the field sampling, and Fabian Deuber and Patrick Baumann for sample preparation, and Annika Ackermann (Grassland Sciences Isolab) for the stable isotope measurements.

## Authors’ Contributions

ADK, KC, NB and A. Gessler conceived the ideas, acquired funding, and designed the methodology. KC carried out the needle and A. Glauser the soil sampling. KC and NB measured the stable isotope traits. A. Glauser measured the starch, sugar and NSC. KC, ADK and A. Glauser analyzed the data, and all authors interpreted the results. KC and ADK wrote the first draft of the manuscript and all authors contributed to the final version.

## Supporting Information

The following Supporting Information is available for this article:

**Table S1:**
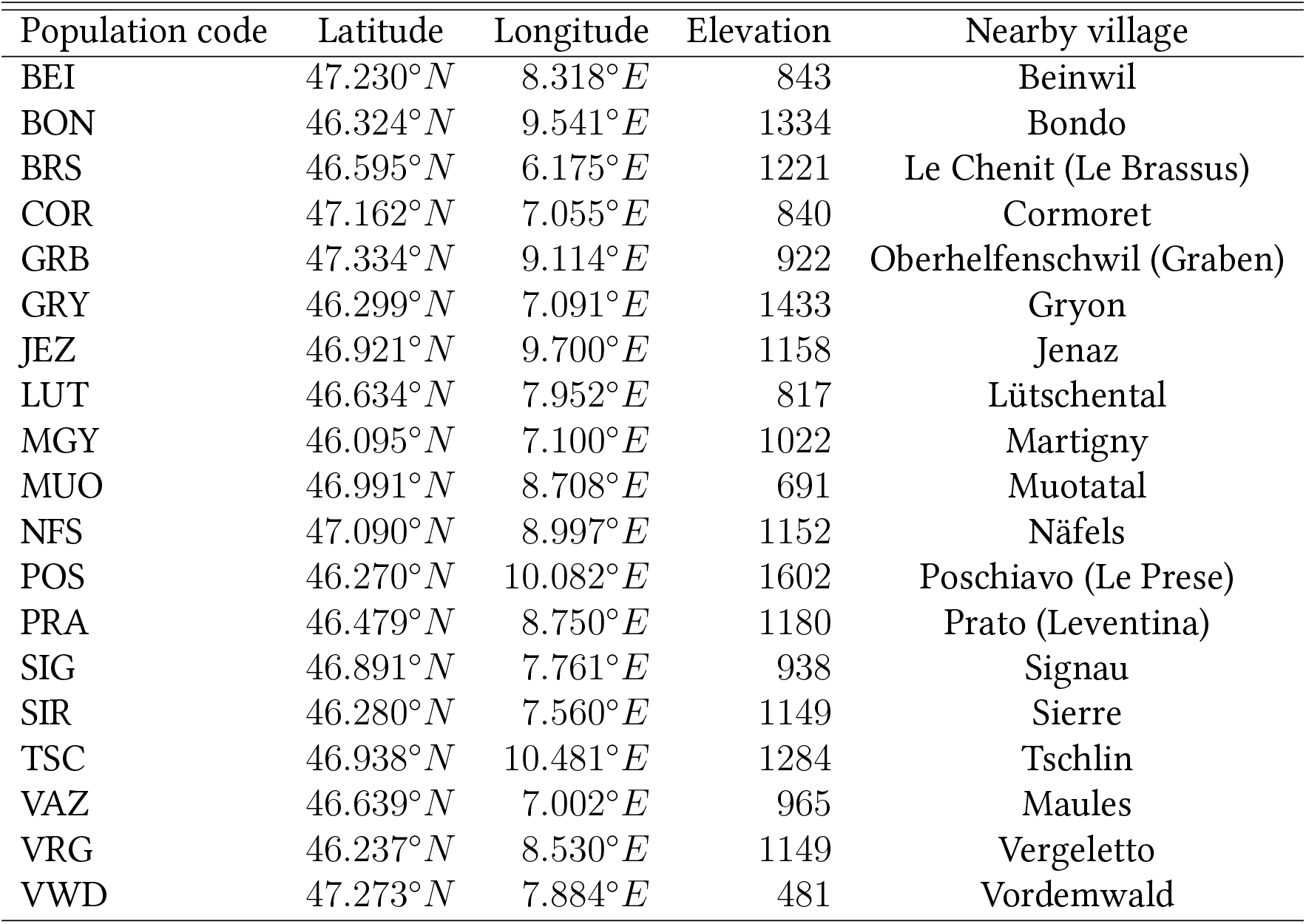
Abbreviations of the population names, their longitude and latitude, elevation in meters, and names of the nearby village (all in Switzerland) from which the abbreviations were derived. Identical to Table S1 of Csilléry *et al*. (2020b).

| Population code | Latitude | Longitude | Elevation | Nearby village |
| --- | --- | --- | --- | --- |
| BEI | 47.230° *N* | 8.318° *E* | 843 | Beinwil |
| BON | 46.324° *N* | 9.541° *E* | 1334 | Bondo |
| BRS | 46.595° *N* | 6.175° *E* | 1221 | Le Chenit (Le Brassus) |
| COR | 47.162° *N* | 7.055° *E* | 840 | Cormoret |
| GRB | 47.334° *N* | 9.114° *E* | 922 | Oberhelfenschwil (Graben) |
| GRY | 46.299° *N* | 7.091° *E* | 1433 | Gryon |
| JEZ | 46.921° *N* | 9.700° *E* | 1158 | Jenaz |
| LUT | 46.634° *N* | 7.952° *E* | 817 | Lütschental |
| MGY | 46.095° *N* | 7.100° *E* | 1022 | Martigny |
| MUO | 46.991° *N* | 8.708° *E* | 691 | Muotatal |
| NFS | 47.090° *N* | 8.997° *E* | 1152 | Näfels |
| POS | 46.270° *N* | 10.082° *E* | 1602 | Poschiavo (Le Prese) |
| PRA | 46.479° *N* | 8.750° *E* | 1180 | Prato (Leventina) |
| SIG | 46.891° *N* | 7.761° *E* | 938 | Signau |
| SIR | 46.280° *N* | 7.560° *E* | 1149 | Sierre |
| TSC | 46.938° *N* | 10.481° *E* | 1284 | Tschlin |
| VAZ | 46.639° *N* | 7.002° *E* | 965 | Maules |
| VRG | 46.237° *N* | 8.530° *E* | 1149 | Vergeletto |
| VWD | 47.273° *N* | 7.884° *E* | 481 | Vordemwald |

**Table S2:** Likelihood ratio tests (LRT) comparing models of different complexity. The full model included all variables listed in the column headings as random effects. This model was compared to a model without the variables in the column headings one by one. If the log likelihood of the full model was greater, the p-value of the LRT is given. p-values less than 0.05 indicate that the model is better including the given random effect. Note however that while the model can be significantly better with a given random effect, the variance component associated with it is not always different from zero (see Fig. 2). If the log likelihood of the reduced model was greater, NA is given.

| **Terminal bud removal** |  |  |  |  |  |  |  |
| --- | --- | --- | --- | --- | --- | --- | --- |
| Year | Trait | Seed Weight | DBH | Treatment | Block | Population | Pedigree |
| **Growth traits** |  |  |  |  |  |  |  |
| 2015 | Height | 0.010 | 0.904 | 0.093 | 0.064 | 0.369 | 0.000 |
|  | Terminal Height | 0.007 | 0.899 | 0.000 | 0.006 | 0.078 | 0.000 |
|  | Diameter | 0.002 | 0.229 | 0.998 | 0.000 | 0.680 | 0.000 |
| 2016 | Height | 0.002 | 0.592 | 0.997 | 0.001 | 0.376 | 0.001 |
|  | Terminal Height | 0.000 | 0.729 | 0.004 | 0.000 | 0.873 | 0.000 |
|  | Diameter | 0.003 | 0.444 | NA | 0.000 | 0.487 | 0.000 |
|  | Fresh weight | 0.003 | 0.502 | 0.999 | 0.000 | 0.544 | 0.000 |
| **Physiological traits** |  |  |  |  |  |  |  |
| 2016 | NSC [%] | 0.496 | 0.386 | 0.888 | 0.055 | 0.104 | 0.103 |
|  | Starch [%] | 0.655 | 0.421 | 0.199 | 0.000 | 0.002 | 0.227 |
|  | Sugar [%] | 0.505 | 0.370 | NA | 0.000 | 0.120 | 0.254 |
|  | C [%] | 0.055 | 0.483 | 0.000 | 0.773 | 0.012 | 0.376 |
| | $\delta^{13}\text{C}$ | 0.278 | 0.526 | NA | 0.000 | 0.298 | 0.145 |
|  | N [%] | 0.524 | 0.255 | 0.106 | 0.095 | 0.007 | NA |
| | $\delta^{15}\text{N}$ | 0.200 | 0.202 | NA | 0.000 | 0.000 | 0.377 |
|  | C/N | 0.301 | 0.386 | 0.057 | 0.153 | 0.028 | NA |
| **Heavy browsing** |  |  |  |  |  |  |  |
| Year | Trait | Seed Weight | DBH | Treatment | Block | Population | Pedigree |
| **Growth traits** |  |  |  |  |  |  |  |
| 2015 | Height | 0.038 | 0.966 | 0.001 | 0.002 | 0.015 | 0.025 |
|  | Terminal Height | 0.111 | 0.765 | 0.000 | 0.001 | 0.004 | 0.099 |
|  | Diameter | 0.005 | 0.325 | 0.997 | 0.000 | 0.889 | 0.000 |
| 2016 | Height | 0.032 | 0.860 | 0.001 | 0.005 | 0.021 | 0.155 |
|  | Terminal Height | 0.011 | 0.796 | 0.000 | 0.002 | 0.076 | 0.107 |
|  | Diameter | 0.005 | 0.493 | 0.095 | 0.000 | 0.846 | 0.000 |
|  | Fresh weight | 0.046 | 0.729 | 0.013 | 0.000 | 0.382 | 0.005 |
| **Physiological traits** |  |  |  |  |  |  |  |
| 2016 | NSC [%] | 0.568 | 0.061 | 0.174 | 0.042 | 0.787 | 0.001 |
|  | Starch [%] | 0.271 | 0.064 | 0.159 | 0.008 | 0.132 | 0.005 |
|  | Sugar [%] | 0.519 | 0.181 | NA | 0.000 | 0.593 | 0.109 |
|  | C [%] | 0.156 | 0.737 | 0.030 | 0.998 | 0.028 | 0.030 |
| | $\delta^{13}\text{C}$ | 0.202 | 0.733 | 0.000 | 0.003 | 0.065 | 0.316 |
|  | N [%] | 0.814 | 0.066 | 0.094 | 0.092 | 0.040 | 0.285 |
| | $\delta^{15}\text{N}$ | 0.559 | 0.563 | 0.000 | 0.002 | 0.432 | 0.073 |
|  | C/N | 0.575 | 0.079 | 0.090 | 0.103 | 0.077 | 0.316 |

**Table S3:** Heritability estimates under control conditions (i.e. no browsing stress) estimated from the pooled model (equation 3) for all traits measured in seedlings in the common garden.

| Trait | $h^2$ | SE |
| --- | --- | --- |
| **Growth traits** |  |  |
| Height 2015 | 0.642 | 0.235 |
| Height 2016 | 0.472 | 0.210 |
| Terminal Height 2015 | 0.375 | 0.169 |
| Terminal Height 2016 | 0.415 | 0.183 |
| Diameter 2015 | 0.656 | 0.413 |
| Diameter 2016 | 0.575 | 0.602 |
| Fresh weight 2016 | 0.586 | 0.237 |
| **Physiology traits (2016)** |  |  |
| NSC [%] | 0.210 | 0.158 |
| Starch [%] | 0.134 | 0.129 |
| Sugar [%] | 0.128 | 0.127 |
| C [%] | 0.094 | 0.124 |
| $\delta^{13}\text{C}$ | 0.184 | 0.150 |
| N [%] | 0.000 | 0.126 |
| $\delta^{15}\text{N}$ | 0.078 | 0.103 |
| C/N | 0.000 | 0.124 |

**Table S4:**
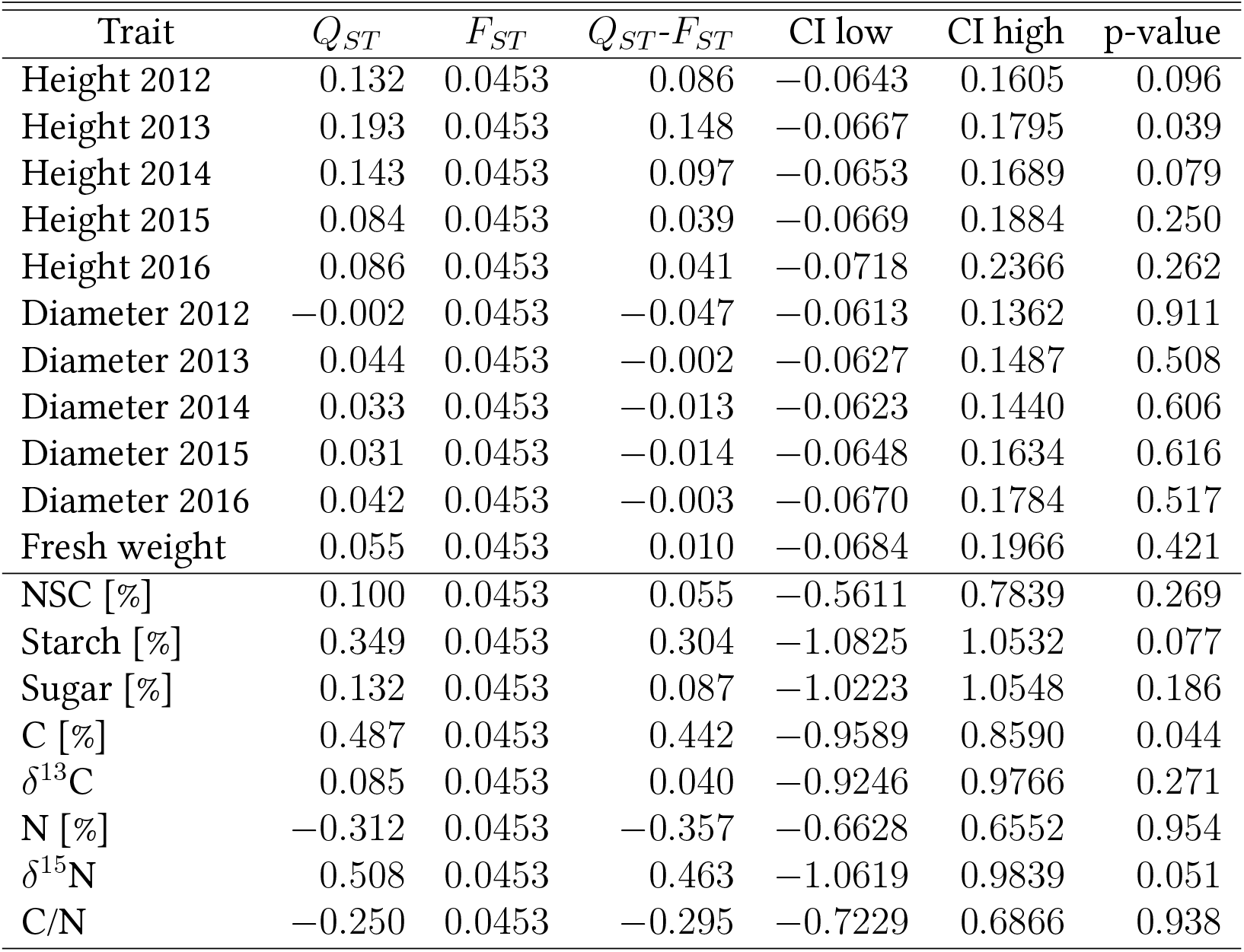
Test of spatially divergence selection based on the comparison between population genetic divergence at traits (*Q*_*ST*_) and genetic divergence at neutral genetic markers (*F*_*ST*_) for all traits measured in seedlings in the common garden. Calculations were performed using the R package *QstFstComp* (Gilbert & Whitlock, 2015). Note that slightly different *Q*_*ST*_ values are reported in Fig. 4 because the calculations are based on a mixed effects model that includes the covariates Seed weight and DBH of mother trees, and random effects block and population. Note also that Csilléry *et al*. (2020b) reported the same test results for 2013 and 2014 Height and Diameter, but their test results were different (higher *Q*_*ST*_ and smaller p-values) because the sample size was larger. The tests reported herein are based on a sample size roughly 40% less due to mortality and because the Heavy browsing blocks are excluded.

**Table S5:**
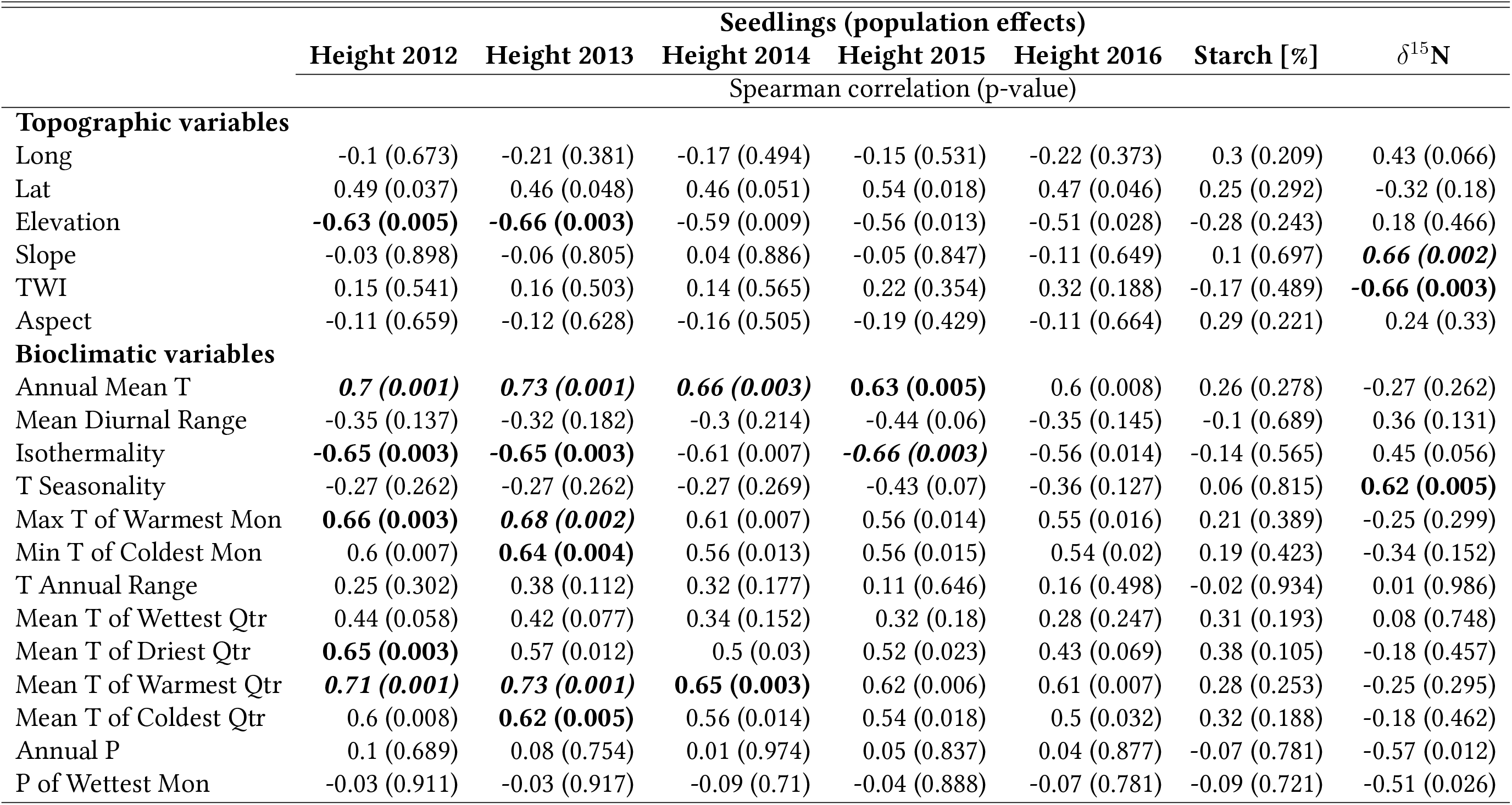
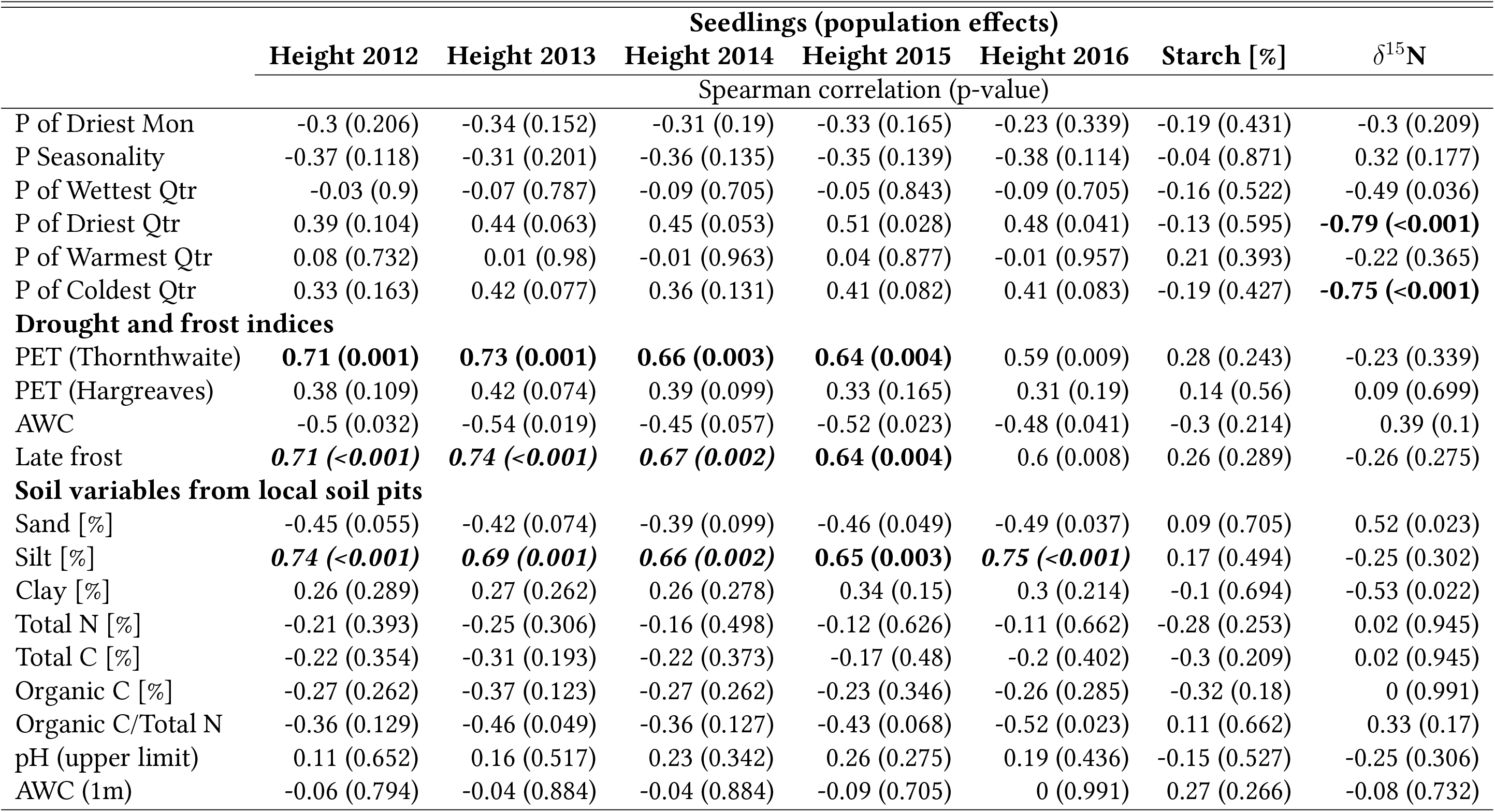
Spearman correlation (uncorrected p-value) between seedling population effects extracted from the pooled model (equation 3) and environmental variables. Correlations that are significant (p-value < 0.05) after a correction for 18 independent tests are marked in ***bold italic***, while those significant with correction for nine independent tests are marked in **bold**. Abbreviations: TWI: topographic wetness index, T: temperature, P: precipitation, Mon: month, Qtr: quarter, PET: potential evapotranspiration, AWC: available water capacity

**Table S6:** Spearman correlation (uncorrected p-value) between traits measured in adult trees *in-situ* (population means) and environmental variables. Correlations that are significant (p-value < 0.05) after a correction for 36 independent tests are marked in ***bold italic***, while those significant with correction for nine independent tests are marked in **bold**. Abbreviations: TWI: topographic wetness index, T: temperature, P: precipitation, Mon: month, Qtr: quarter, PET: potential evapotranspiration, AWC: available water capacity

|  | Adult trees in-situ (population median) |  |  |  |  |
| --- | --- | --- | --- | --- | --- |
| | C [%] | $\delta^{13}\text{C}$ | N [%] | $\delta^{15}\text{N}$ | C/N |
|  | Spearman correlation (p-value) |  |  |  |  |
| Topographic variables |  |  |  |  |  |
| Longitude | -0.6 (0.008) | -0.13 (0.604) | -0.38 (0.108) | 0.11 (0.657) | 0.32 (0.182) |
| Latitude | -0.14 (0.58) | -0.33 (0.163) | 0.33 (0.161) | -0.01 (0.957) | -0.34 (0.15) |
| Elevation | -0.02 (0.94) | 0.45 (0.051) | -0.15 (0.529) | 0.18 (0.471) | 0.16 (0.508) |
| Slope | -0.49 (0.034) | -0.05 (0.839) | **-0.81 (<0.001)** | -0.1 (0.683) | **0.78 (<0.001)** |
| TWI | 0.47 (0.045) | -0.11 (0.665) | **0.61 (0.005)** | -0.25 (0.295) | -0.56 (0.015) |
| Aspect | 0.46 (0.049) | 0.34 (0.15) | -0.12 (0.617) | -0.27 (0.264) | 0.12 (0.623) |
| Bioclimatic variables |  |  |  |  |  |
| Annual Mean T | 0.02 (0.945) | -0.4 (0.087) | 0.16 (0.522) | -0.2 (0.402) | -0.17 (0.48) |
| Mean Diurnal Range | -0.12 (0.621) | 0.18 (0.466) | -0.46 (0.047) | -0.19 (0.423) | 0.42 (0.073) |
| Isothermality | -0.1 (0.689) | 0.29 (0.228) | -0.31 (0.201) | -0.06 (0.809) | 0.3 (0.211) |
| T Seasonality | -0.28 (0.247) | 0.04 (0.87) | **-0.65 (0.003)** | -0.07 (0.776) | 0.59 (0.009) |
| Max T of Warmest Mon | 0.11 (0.667) | -0.37 (0.116) | 0.13 (0.61) | -0.21 (0.385) | -0.12 (0.631) |
| Min T of Coldest Mon | 0.17 (0.48) | -0.33 (0.17) | 0.26 (0.288) | -0.19 (0.444) | -0.24 (0.324) |
| T Annual Range | 0.06 (0.809) | -0.28 (0.244) | -0.23 (0.351) | -0.41 (0.082) | 0.17 (0.48) |
| Mean T of Wettest Qtr | -0.22 (0.358) | -0.59 (0.008) | 0.11 (0.643) | -0.3 (0.217) | -0.1 (0.689) |
| Mean T of Driest Qtr | -0.24 (0.313) | -0.34 (0.148) | -0.04 (0.869) | -0.06 (0.809) | -0.03 (0.9) |
| Mean T of Warmest Qtr | 0.03 (0.905) | -0.37 (0.119) | 0.1 (0.675) | -0.22 (0.369) | -0.11 (0.641) |
| Mean T of Coldest Qtr | 0.12 (0.631) | -0.38 (0.113) | 0.13 (0.605) | -0.26 (0.272) | -0.11 (0.641) |
| Annual P | 0.04 (0.871) | -0.12 (0.637) | 0.52 (0.022) | 0.16 (0.503) | -0.6 (0.008) |
| P of Wettest Mon | 0.11 (0.667) | -0.08 (0.748) | 0.58 (0.009) | 0.12 (0.616) | **-0.64 (0.004)** |
| P of Driest Mon | **0.63 (0.005)** | 0.09 (0.705) | 0.08 (0.747) | -0.25 (0.295) | -0.01 (0.968) |
| P Seasonality | -0.04 (0.865) | -0.07 (0.762) | 0.47 (0.043) | 0.05 (0.854) | -0.43 (0.068) |
| P of Wettest Qtr | 0.08 (0.748) | -0.07 (0.778) | 0.46 (0.048) | 0.26 (0.285) | -0.51 (0.027) |
| P of Driest Qtr | 0.28 (0.24) | -0.04 (0.878) | 0.46 (0.048) | 0.02 (0.928) | -0.47 (0.043) |
| P of Warmest Qtr | -0.2 (0.414) | 0.02 (0.943) | 0.21 (0.389) | 0.09 (0.705) | -0.29 (0.231) |
| P of Coldest Qtr | 0.18 (0.457) | -0.1 (0.692) | 0.53 (0.018) | 0.01 (0.968) | -0.57 (0.012) |
| Drought and frost indices |  |  |  |  |  |
| PET (Thornthwaite) | -0.04 (0.877) | -0.43 (0.064) | 0.1 (0.68) | -0.19 (0.431) | -0.11 (0.641) |
| PET (Hargreaves) | 0.19 (0.431) | -0.26 (0.286) | -0.2 (0.42) | -0.35 (0.145) | 0.25 (0.299) |
| late.frost2 | -0.01 (0.974) | -0.44 (0.061) | 0.13 (0.597) | -0.19 (0.427) | -0.14 (0.556) |
| Soil variables from local soil pits |  |  |  |  |  |
| Sand [%] | -0.06 (0.815) | 0.13 (0.584) | -0.18 (0.455) | 0.18 (0.471) | 0.2 (0.402) |
| Silt [%] | -0.11 (0.652) | -0.1 (0.684) | -0.1 (0.691) | -0.35 (0.147) | 0.06 (0.804) |
| Clay [%] | -0.12 (0.616) | -0.13 (0.596) | 0.25 (0.296) | -0.12 (0.61) | -0.3 (0.217) |
| Total N [%] | 0.12 (0.631) | -0.04 (0.881) | 0.11 (0.656) | 0.25 (0.306) | -0.12 (0.636) |
| Total C [%] | 0.1 (0.683) | -0.06 (0.792) | 0.08 (0.755) | 0.36 (0.133) | -0.06 (0.809) |
| Total Organic C [%] | 0.09 (0.71) | -0.09 (0.726) | 0.04 (0.864) | 0.33 (0.163) | -0.03 (0.9) |
| Organic C/Total N | -0.13 (0.595) | -0.02 (0.935) | -0.58 (0.009) | 0.03 (0.911) | **0.63 (0.005)** |
| pH (upper limit) | -0.04 (0.888) | 0.08 (0.756) | 0.06 (0.799) | 0.08 (0.754) | -0.07 (0.776) |
| AWC (1m) | 0.02 (0.935) | -0.02 (0.949) | 0.04 (0.855) | -0.57 (0.011) | 0 (0.991) |

**Figure S1:**
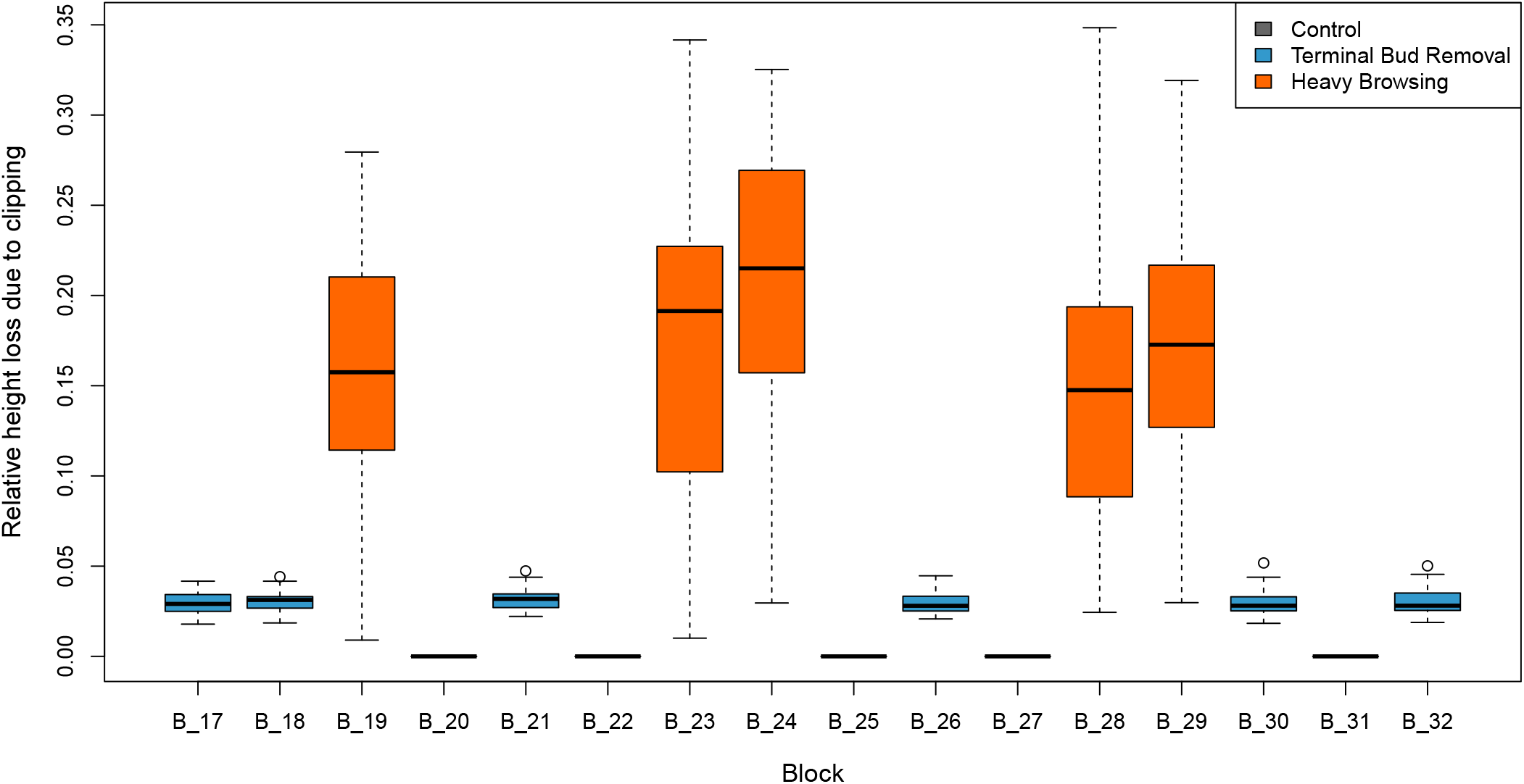
Height loss per block. See Fig/ 1 for the spatial arrangement of the blocks.

**Figure S2:**
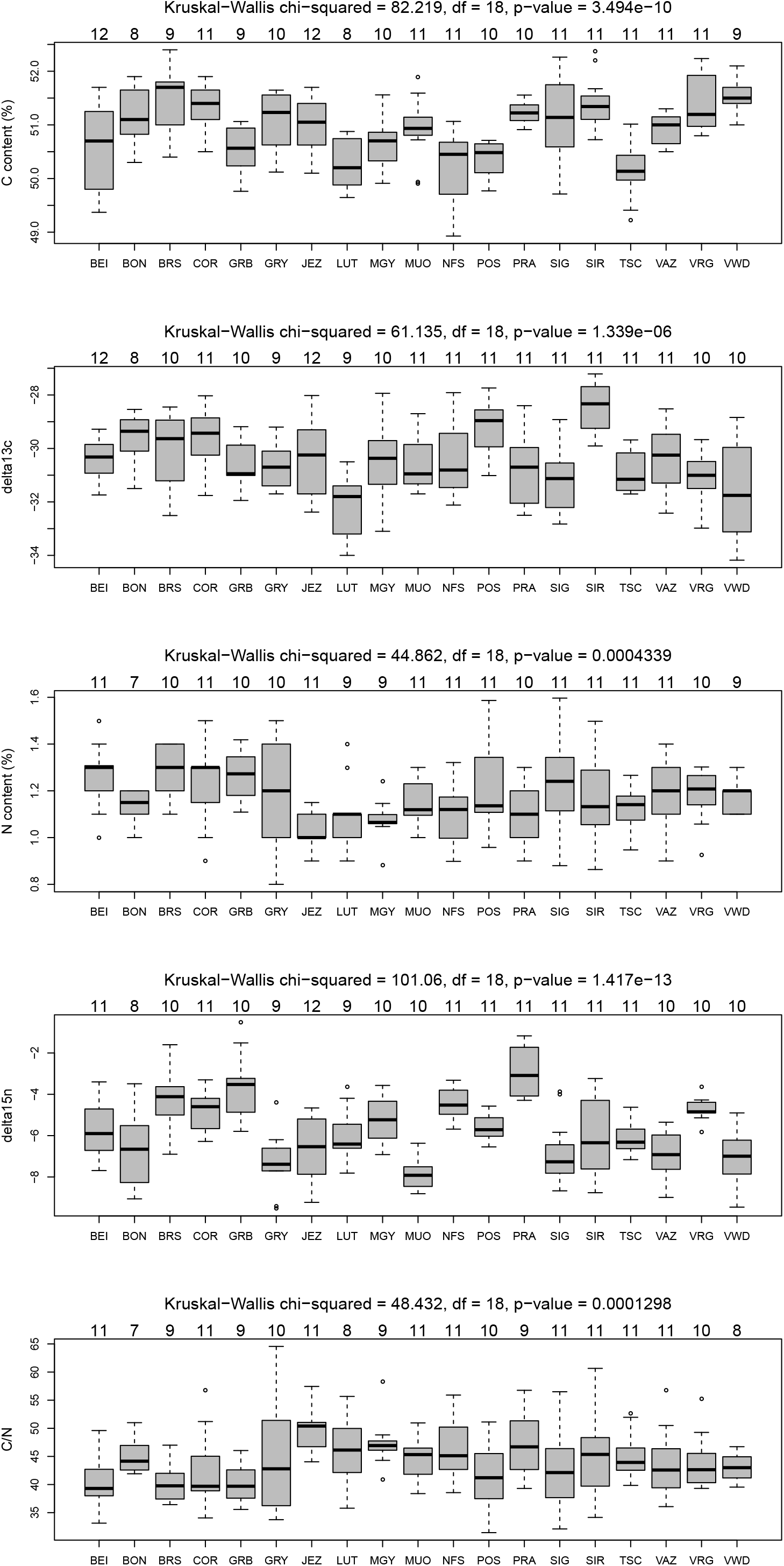
Traits measured in adult trees *in-situ* across the 19 populations and Kruskal-Wallis test of differences between population medians1.0

**Figure S3:**
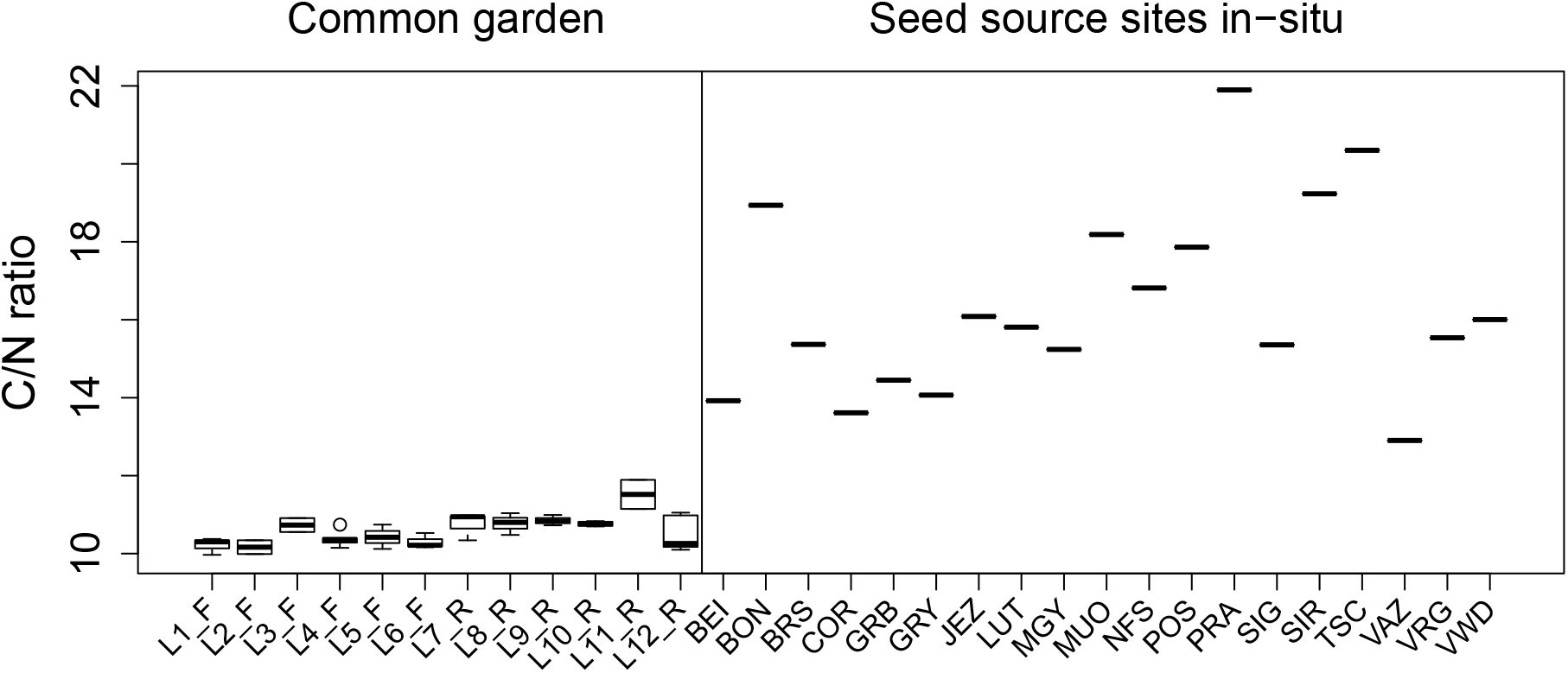
C/N ratio of the top mineral soil layer at the common garden site in Matzendorf and at the seed source populations. Boxplots at the common garden site represent variation of C/N across three to four replicate measures between layers 0 to 15 cm depth. A single measure was taken in-situ.

**Figure S4:**
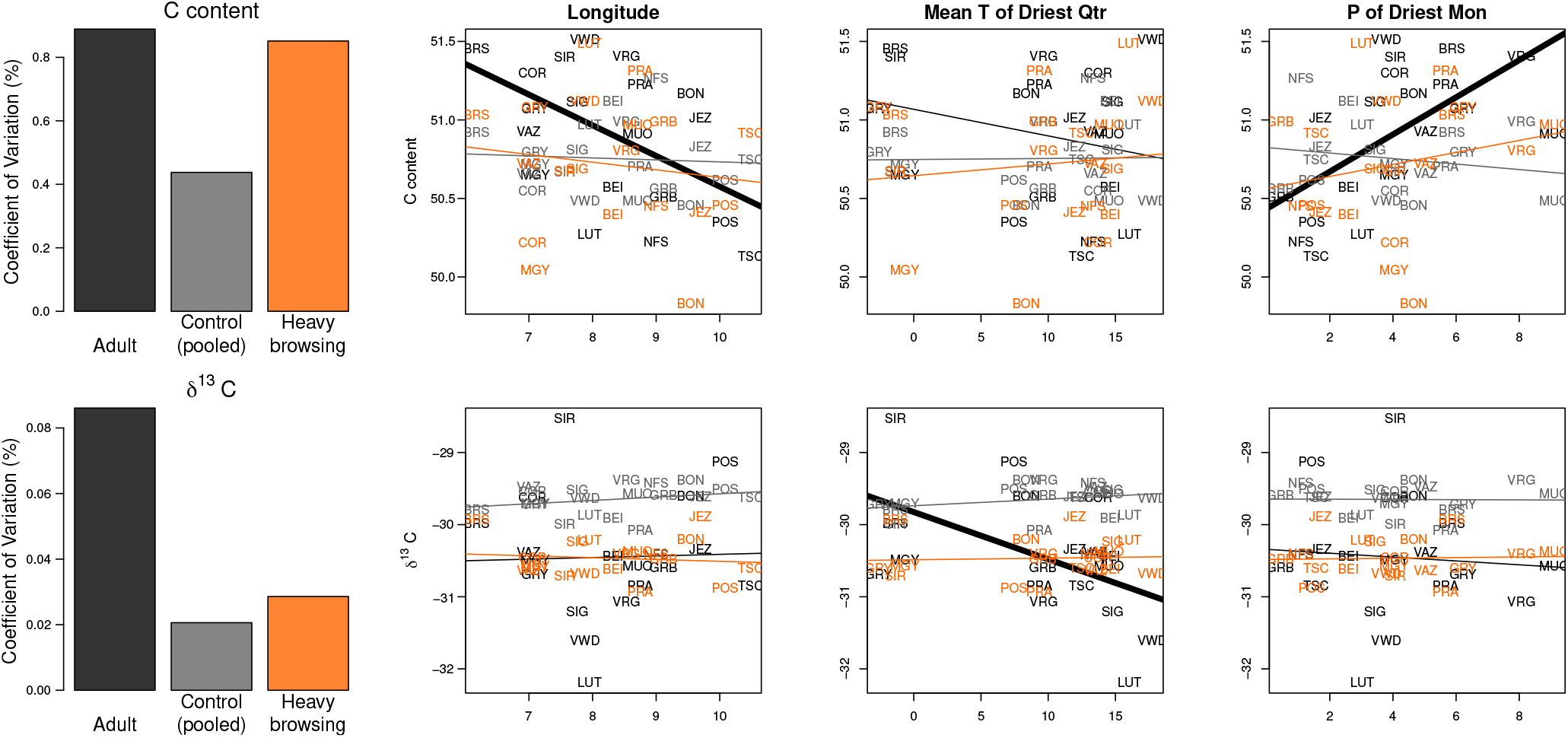
Coefficient of variation in C concentration and *δ*^13^*C*, and correlation between adult population medians (and seedling population effects) and environmental variables. Note that these are the strongest correlations that have unadjusted p-values < 0.05 (see Table S6), but none of them passed the correction for multiple testing.

